# Loss of CHCHD2 and CHCHD10 activates OMA1 peptidase to disrupt mitochondrial cristae phenocopying patient mutations *in vivo*

**DOI:** 10.1101/773135

**Authors:** Yi-Ting Liu, Xiaoping Huang, Diana Nguyen, Eszter Dombi, Danielle A Springer, Joanna Poulton, Shiori Sekine, Derek P Narendra

**Author notes:** Joint first authors/equal contributing authors (Yi-Ting Liu, Xiaoping Huang). To whom correspondence should be addressed. Address: 35 Convent Drive, Bldg 35 Rm 2A215, Bethesda, MD, 20892.

## Abstract

Dominant mutations in the mitochondrial paralogs CHCHD2 (C2) and CHCHD10 (C10) were recently identified as causing Parkinson’s disease and ALS/FTD/myopathy, respectively. Disruption of mitochondrial cristae has been observed in mutant C10 patient tissues and animal models, but the mechanism for this disruption remains controversial. Additionally, C10 patient mutant knock-in (KI) mice were recently reported to activate a mitochondrial integrated stress response (mt-ISR) and develop cardiomyopathy not seen in C10 knockout (KO) mice, calling into question whether mutant C10 pathogenesis is related to C2/C10 normal function or purely toxic gain of function. Here, using the first C2/10 double knockout (DKO) mice, we report that C10 pathogenesis and the normal function of C2/10 are intimately linked. Similar to patients with C10 mutations, we found that C2/10 DKO mice (but not either single KO mice) have disrupted mitochondrial cristae, due to cleavage of the mitochondrial shaping protein L-OPA1 by the stress-induced peptidase OMA1. OMA1 was found to be activated similarly in affected tissues of mutant C10 KI mice, demonstrating that L-OPA1 cleavage is a novel mechanism for cristae abnormalities due to both C10 mutation and C2/C10 loss, and that OMA1, a driver of neurodegeneration in other contexts, may be a therapeutic target. Finally, C2/10 DKO mice partially phenocopied mutant C10 KI mice with the development of cardiomyopathy and activation of the mt-ISR in affected tissues, tying mutant C10 pathogenesis to C2/C10 function.

## Introduction

Autosomal dominant mutations in the mitochondrial paralogs C2 and C10 were recently shown to cause, respectively, Parkinson’s disease (PD) and a spectrum of neuromuscular disorders (that includes myopathy and Amyotrophic Lateral Sclerosis [ALS]) (Bannwarth et al., 2014; Funayama et al., 2015; Ajroud-Driss et al., 2015; Penttilä et al., 2015). Most mutations with proven pathogenicity in C2/C10 cluster in a common hydrophobic α-helix N-terminal to the coiled-helix-coiled-helix (CHCH) domain, consisting of dual CX_9_C motifs. These include the C10 S59L substitution, which was originally identified in a family with ALS, frontotemporal dementia, and myopathy, and substitution of the neighboring residue, G58R, identified in a family with myopathy (Bannwarth et al., 2014; Ajroud-Driss et al., 2015).

C2 and C10 are co-expressed in relevant cellular subtypes, such as *substantia nigral* neurons, pyramidal neurons, and myocytes, where they directly interact within a ∼200 kDa complex in the mitochondrial intermembrane space (Straub et al., 2018; Burstein et al., 2018; Huang et al., 2018). Although their precise function remains unknown, cristae abnormalities have been observed in muscle tissue and cell lines from patients with C10 mutations, as well as a *Drosophila* lacking the C2/C10 ortholog (Bannwarth et al., 2014; Ajroud-Driss et al., 2015; Meng et al., 2017). To explain cristae abnormalities in C2/C10 patients, C10 and later C2 were proposed to be structural components of the mitochondrial contact site and cristae organizing system (MICOS) complex (Bannwarth et al., 2014; Genin et al., 2016; Zhou et al., 2019). However, recent reports have challenged this model, suggesting that C2/C10 mutation or loss may lead to cristae abnormalities by another as of yet unidentified mechanism (Straub et al., 2018; Burstein et al., 2018; Huang et al., 2018).

Cristae are infoldings of the inner mitochondrial membrane and can be considered the fundamental bioenergetic unit within mitochondria. Studded with respiratory complexes and rimmed with dimers of the F_1_F_o_ ATP synthase, they increase surface area for oxidative phosphorylation (Davies et al., 2011). Cristae are shaped by at least three protein complexes: the MICOS complex, bending the cristae membrane at its junction with the boundary membrane; the GTPase OPA1, which mediates inner membrane fusion and supports cristae structure by bridging apposing membranes in the cristae fold; and ATP synthase dimers, which bend the membrane at the cristae edge (Paumard et al., 2002; von der Malsburg et al., 2011; Harner et al., 2011; Hoppins et al., 2011; Frezza et al., 2006; Dudkina et al., 2006; Davies et al., 2012; Ban et al., 2017).

OPA1, in particular, is highly regulated by mitochondrial bioenergetics and proteostatic stress to dynamically shape the inner membrane in response to changing conditions within the mitochondrial reticulum. The peptidase OMA1 is activated by mitochondrial stressors to cleave the active long form of OPA1 (L-OPA1) from its membrane anchor, leading to mitochondrial fragmentation and alterations in cristae structure (Head et al., 2009; Ehses et al., 2009). L-OPA1 processing by OMA1 also occurs upon loss of quality control proteases, such as YME1, AFG3L2, and SPG7, as well as inner membrane scaffolding proteins of the SPFH family, namely, prohibitins and SLP-2, that organize quality control proteases within inner membrane microdomains (Anand et al., 2014; Wai et al., 2016; Merkwirth et al., 2008; Ehses et al., 2009; Magri et al., 2018). Thus, OMA1 provides an escape mechanism for cristae with failed of protein quality control, separating dysfunctional mitochondrial units from the reticulum for degradation by autophagy (MacVicar and Lane, 2014).

Very recently, mice null for C10 and knock-in for the pathogenic S59L mutation have been reported (Genin et al., 2019; Anderson et al., 2019; Burstein et al., 2018). Whereas C10^-/-^ mice have a normal lifespan, C10^S59L/+^ knock-in (KI) mice develop a cardiomyopathy by 23 weeks, which is uniformly fatal at 13 months, supporting a gain of function mechanism of pathogenesis. Anderson et al. (2019), in particular, observed C10 levels increase and co-aggregate with C2 in affected tissues, leading them to propose a toxic aggregate model of pathogenesis, in which C2/C10 toxic aggregates trigger the integrated mitochondrial stress response (mt-ISR) and tissue degeneration. However, it remains unclear in this aggregate model whether the pathogenesis of mutant C10 relates to C2/C10 physiologic function.

Here, we evaluate the first C2/C10 DKO mouse model with two novel findings. First, we identify OPA1 cleavage by the stress-induced peptidase OMA1 to be at least partially responsible for cristae abnormalities in absence of C2/C10. We further show that OMA1 activation is a key event in mutant C10 pathogenesis *in vivo*. Thus, we establish for the first time OPA1 cleavage as potential mechanism for cristate abnormalities in patients with C10 mutations and C2/C10 models. Second, we observe that C2/10 DKO mice unlike C10 KO mice partially phenocopies the recently reported mutant C10 model, including activation of the mt-ISR and development of cardiomyopathy. This challenges at least the simplest version of the C10 aggregate model of pathogenesis, in which misfolded C10 protein directly activates mt-ISR and causes tissue degeneration irrespective of its normal function; rather, our findings suggest that C10 pathogenesis and C2/10 physiological function are intimately linked.

## Results

### C2/C10 double knockout leads to cristae abnormalities due to increased L-OPA1 processing by the stress-induced protease OMA1 in cell culture

To investigate the physiological role of C2 and C10 in mammalian primary cells, we generated C2 and C10 single knockout (KO) and double knockout (DKO) mice (Figure 1A). Whereas cristae structure was normal for C2 and C10 single knockout cell lines by Transmission Electron Microscopy (TEM) (Supplemental Figure 1A), primary fibroblasts from DKO mice exhibited abnormalities in mitochondrial ultrastructure, including convoluted cristae, which were sometimes circular and detached from boundary membrane, as well as a decrease in cristae number (6.778 vs. 14.46 cristae/μm^2^, p < 0.0001) (Figure 1B – D).

**Figure 1.**
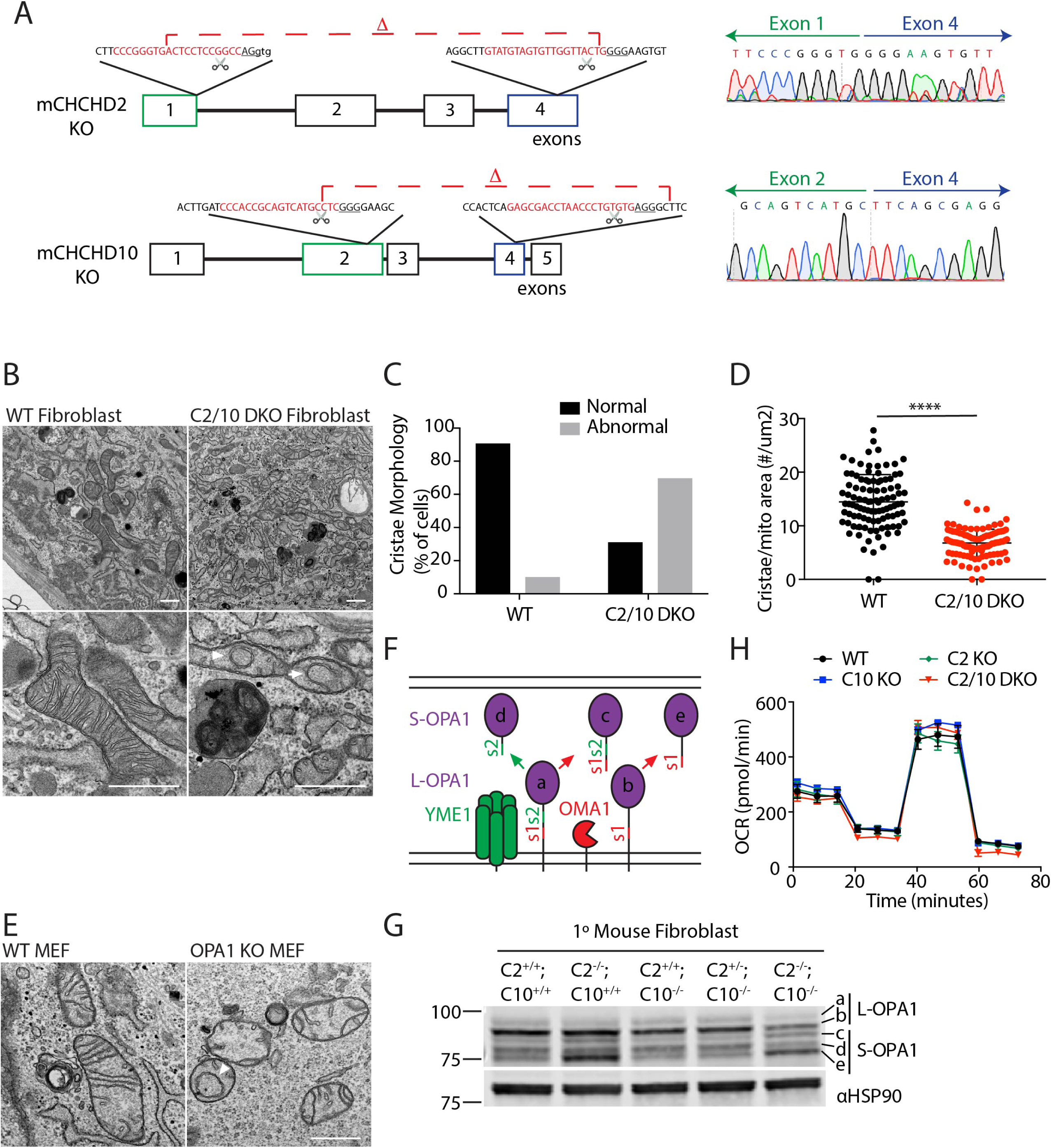
Cristae abnormalities in C2/C10 DKO primary fibroblasts associated with OPA1 processing by OMA1 activation. (A) C2 and C10 KO mice were generated by CRIPSR/Cas9 genomic editing with CRISPR cut sites (scissor icon) and guide RNA sequence (red text) and PAM sequence (underlined) indicated. Sanger sequencing across the deletions verifies loss of the intervening exons. (B) Representative TEM images of WT and C2/C10 DKO fibroblast cells. Arrowhead indicates detached cristae and arrow points to convoluted cristae. (C) Quantification of abnormal mitochondria for > 100 mitochondria per condition. (D) The number of cristae per mitochondrion area (μm^2^) was calculated for > 100 mitochondria per condition. (E) Representative TEM images of transformed WT and OPA1 KO MEF cells. Arrowhead indicates detached cristae. (F) Model depicts processing of the two membrane-bound long forms of OPA1 (L-OPA1) at protease sites s1 and s2 into the three short forms (S-OPA1) by the proteases OMA1 (at the s1 site) and YME1 (at the s2 site). (G) Immunoblot from lysates of WT, C2^-/-^, C10^-/-^, C2^+/-^;C10^-/-^, and C2^-/-^;C10^-/-^ (DKO) primary mouse fibroblasts. Immunoblot is representative of 3 biological replicates. HSP90 serves as the loading control. OPA1 isoforms (a – e) depicted in (F) are indicated. (H) Oxygen consumption rates (OCR) of WT, C2^-/-^, C10^-/-^, C2^-/-^;C10^-/-^ (DKO) primary fibroblasts measured by the Seahorse instrument. Scale bars in all images = 600 nm.

As the cristae abnormalities in C2/C10 DKO fibroblasts resembled those with OPA1 KO (Figure 1E), we assessed OPA1 processing by immunoblotting. There are five OPA1 isoforms visible by immunoblotting corresponding to the membrane-bound long OPA1 (L-OPA1) forms (*a* and *b*), and their proteolytic cleavage products, the short OPA1 (S-OPA1) forms (*c* – *e*), generated by YME1 (*d*) or OMA1 cleavage (*c* and *e*), respectively (Figure 1F) (Ehses et al., 2009; Head et al., 2009). In C2/C10 DKO fibroblasts, L-OPA1 forms decreased and OMA1-dependent S-OPA1 forms (*c* and *e*) increased, consistent with OMA1 cleavage (Figure 1G). Notably, L-OPA1 processing was intermediate in C2 single KO cells and normal in C10 KO cells, consistent with partial functional redundancy of the paralogs.

In contrast to L-OPA1 levels, DRP1 and Mitofusin-2 levels were similar in all cell lines (Supplemental Figure 1B). Additionally, protein levels of the MICOS subunits, Mic60, Mic27, Mic25, and Mic19, were similar in C10 KO fibroblasts, which did not have cristae abnormalities, and C2/C10 DKO fibroblasts, which had cristae abnormalities, from litter matched animals (Figure 1B and Supplemental Figure 1A and B). Levels of MICOS subunits in both C10 KO and C2/C10 DKO fibroblasts were reduced when compared to a WT fibroblast lines from unrelated mice (Supplemental Figure 1B). This suggested either that loss of C10 leads to decreased MICOS subunits but not cristae defects or that there is variability of MICOS protein levels in fibroblast lines from unrelated mice. In either case, these results indicated that a decrease in MICOS levels is not responsible for the cristae defects observed in C2/10 DKO mouse fibroblasts.

As OMA1 is activated in response to bioenergetic collapse, we investigated cellular respiration in DKO cells by oximetry. Basal and maximal oxygen consumption was comparable for DKO and WT primary fibroblasts under growth conditions in which L-OPA1 processing was disrupted (Figure 1H). Mitochondrial membrane potential measured by TMRM fluorescence was similar for C10 KO and C2/10 DKO fibroblasts (Supplemental Figure 1C). Membrane potential in C2/C10 single and double knockouts was slightly reduced when compared to WT lines (Supplemental Figure 1C). Together these findings argue against OPA1 instability in C2/10 DKO cells resulting from disrupted mitochondrial bioenergetics.

To assess whether OPA1 processing was altered also in human cells, we evaluated the OMA1/OPA1 axis in HEK293 DKO cells that we described previously (Huang et al., 2018). Similar to DKO mouse fibroblasts, cristae number was significantly reduced in HEK293 DKO cells (Figure 2A and B). Likewise, L-OPA1 was found to be processed in a pattern reflective of OMA1 activation (Figure 2C). OMA1 levels were likewise decreased, as expected given OMA1 is degraded shortly after its activation (Head et al., 2009). Similar to primary fibroblasts we previously found only a mild oxygen consumption deficit in HEK293 DKO cells compared to wildtype cells (Huang et al., 2018), and TMRE fluorescence was slightly higher for DKO cells compared to wildtype cells (Supplemental Figure 1D). Processing of another substrate of activated OMA1, PGAM5, was unaltered in DKO cells, suggesting that OMA1 activation with C2/C10 DKO may preferentially affect L-OPA1 processing (Supplemental Figure 2A) (Sekine et al., 2012). To formally test whether OMA1 is responsible for L-OPA1 cleavage in the absence of C2/C10, we generated a C2/C10/OMA1 triple KO (TKO) line. L-OPA1 isoforms (*a* and *b*) were stabilized against cleavage to isoforms *c* and *e*, whereas basal processing of L-OPA1 isoform *a* to *d* by the protease YME1 was unaffected (Figure 2C). Importantly, cristae density was partially restored in TKO cells (Figure 2B). Together these findings demonstrate cristae abnormalities in the absence of C2 and C10 are caused at least in part by OMA1 cleavage of L-OPA1.

**Figure 2:**
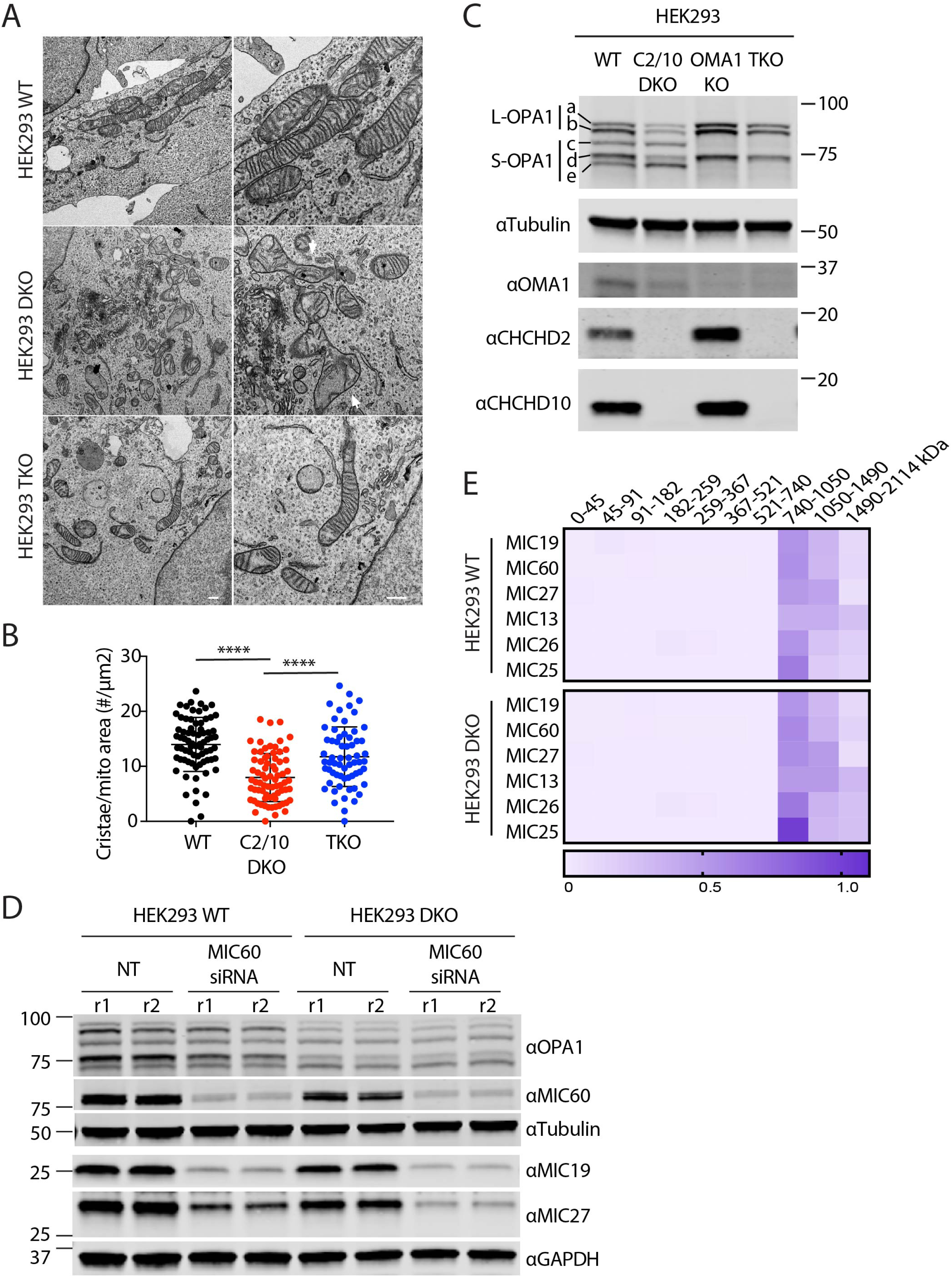
Cristae abnormalities in C2/C10 DKO cells are due at least in part to OMA1 activation. (A) Representative TEM images of WT, C2/C10 DKO, and C2/C10/OMA1 triple KO (TKO) HEK293 cells. Scale bar = 800 nm. Arrows indicate abnormal mitochondria. (B) The number of cristae per mitochondrion area (μm^2^) was calculated for > 100 mitochondria per condition. (C) Immunoblot from lysates of WT, C2/C10 DKO, OMA1 KO, and TKO HEK293 cells. Tubulin served as the loading control. (D) Immunoblot of WT and C2/C10 DKO HEK293 cells following knockdown of MIC60 or left untransfected (NT). Immunoblot is representative of 3 biological replicates obtained on 2 different occasions. Lysates from 2 replicates (r1 and r2) are shown. (E) Heat map depicts relative abundance of MICOS complex subunits in SILAC labelled WT and DKO HEK293 mitochondria, solubilized in 1% digitonin and separated by BN-PAGE prior to detection by quantitative mass spectrometry. Columns in the heat map represent gel slices at the indicated position. Rows indicate proteins detected.

As the MICOS complex is reported to interact with OPA1 and with C2/C10 (Ding et al., 2015; Darshi et al., 2011; Genin et al., 2016; Zhou et al., 2019), we tested whether reduction in MICOS subunits could account for the observed OMA1 activation and L-OPA1 cleavage. As expected, depletion of a central subunit of the MICOS complex, MIC60, destabilized MIC19 and MIC27, components of the respective MICOS subcomplexes (Figure 2D). OPA1 processing, by contrast, was unaffected under these conditions, demonstrating that MICOS complex instability alone cannot account for OPA1 cleavage by OMA1.

We additionally assessed the relative abundance of the MICOS and other complexes previously found to affect OMA1 activation and OPA1 cleavage by LC-MS/MS-based complexomics in WT and DKO HEK293 cells. Detected subunits of the SPY, Prohibitin, and MICOS complexes were of similar abundance in WT and DKO cells (Figure 2E, Supplemental Figure 2B, and Table S1).

Finally, we assessed for interactions among C2, OPA1, OMA1, and proteins forming the SPY complex, SLP-2, PARL, and YME1 (Wai et al., 2016). As reported, previously, C10 robustly co-immunoprecipitates with C2-Flag (Supplemental Figure 2C and D). By contrast, OPA1 and OMA1 failed to co-immunoprecipitate with C2 under the same conditions. Interactions between C2 and PARL and C10 and YME1 of the SPY complex have been identified, previously, in LC-MS/MS co-immunoprecipitation experiments (Floyd et al., 2016; Wai et al., 2016). We were able to confirm an interaction between C2-Flag and YME1. However, we were not able to detect interactions between C2-Flag and PARL or C2-Flag and SLP-2 under the conditions used (Supplemental Figure 2C and D). Similarly, whereas PARL-Flag pulled down endogenous SLP-2, it failed to pull down endogenous C2 or C10. Together these findings could be consistent with weak transient interactions with the SPY complex but argue against strong physical interactions between C2 and OMA1, the SPY complex, or OPA1.

### Mitochondrial fragmentation by C10 mutants depends on L-OPA1 cleavage by OMA1

Having identified a functional interaction between C2/C10 and OMA1/OPA1, we next asked whether OMA1 activation is responsible for mitochondrial fragmentation that we and others observed previously following overexpression of the pathogenic C10 mutation G58R in human cells (Ajroud-Driss et al., 2015; Huang et al., 2018). Over-expression of C10 G58R caused severe mitochondrial fragmentation on a wildtype background, as was expected (Figure 3A and B). Notably, this fragmentation was blocked by OMA1 KO. Milder fragmentation observed with over-expression of wildtype C10 was similarly blocked in OMA1 KO cells (Figure 3A and B). Consistently, C10 G58R over-expression strongly stimulated processing of L-OPA1 to S-OPA1 whereas wildtype C10 had a weaker effect; and both were blocked by OMA1 KO (Figure 3C). These findings demonstrated that C10 G58R overexpression causes mitochondrial fragmentation by triggering cleavage of L-OPA1 by OMA1.

**Figure 3:**
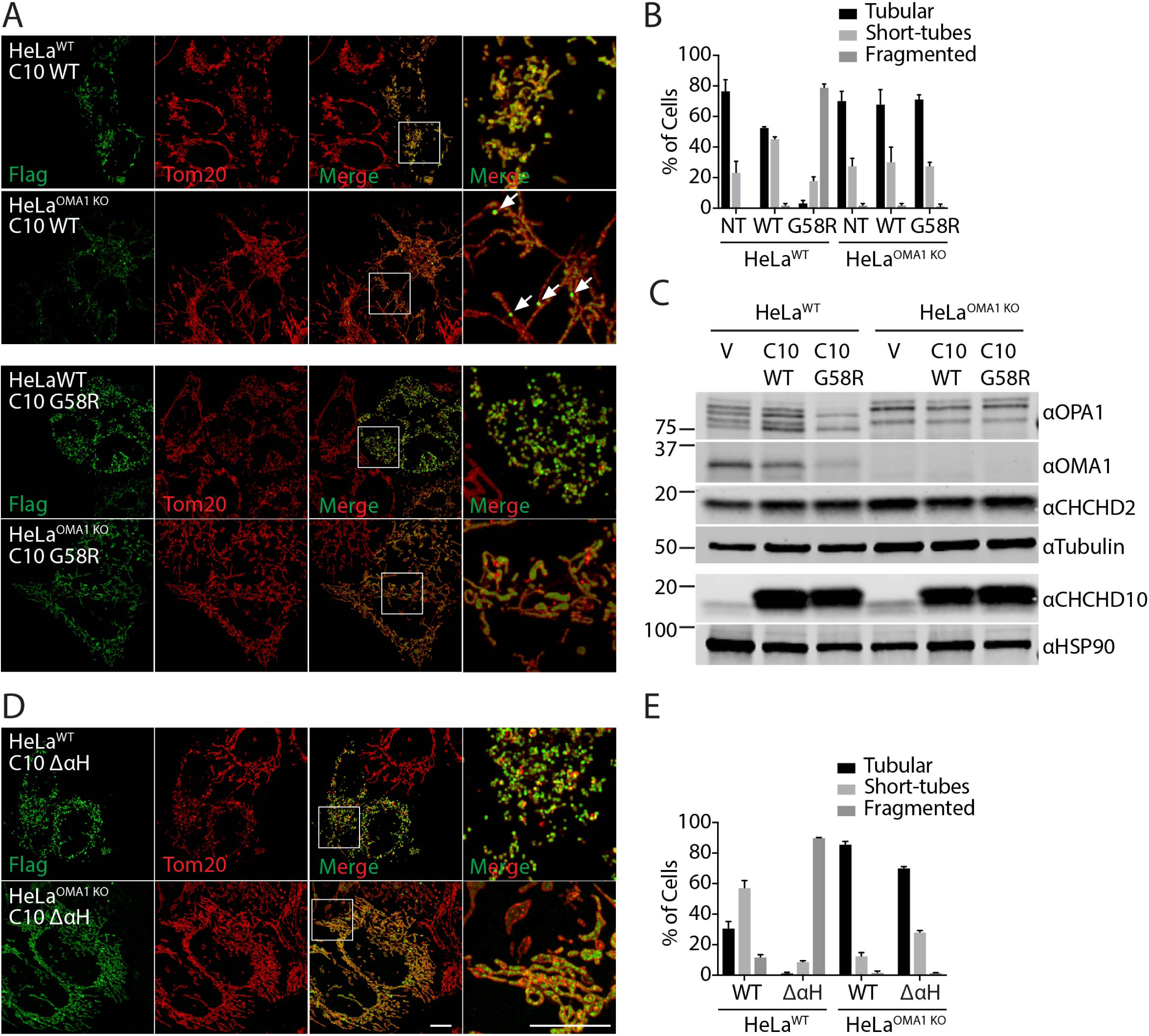
Mitochondrial fragmentation induced by C10 mutant depends on OMA1 processing of OPA1. (A) Representative Airyscan confocal images of WT and OMA1 KO HeLa cells transiently transfected with C10 WT-Flag (C10 WT) or C10 G58R-Flag (C10 G58R). Anti-Flag immunostaining in green and anti-Tom20 in red. (B) Quantification of mitochondrial morphology depicted in (A) for > 150 cells in 3 biological replicates per condition. Untransfected cells (NT) were identified by the absence of Flag staining in each of the samples. Counts were pooled as they did not differ among the samples. (C) Immunoblot from lysates of WT and OMA1 KO HeLa cells transduced with vector (V), C10 WT (C10 WT), or C10 G58R (C10 G58R). Tubulin and HSP90 serve as the loading controls in the respective blots. Immunoblot is representative of 3 biological replicates. (D) Representative Airyscan confocal images of WT and OMA1 KO HeLa cells transiently transfected with C10 WT-Flag (C10 WT) or C10-Flag with the N-terminal α-helix containing G58 deleted (C10 ΔαH). Anti-Flag immunostaining in green and anti-Tom20 in red. (E) Quantification of mitochondrial morphology in (A) for > 150 cells in 3 biological replicates per condition. Scale bars in all images = 10 μm. Except where indicated all confocal images were acquired as a z-stack and represented as a maximum projection image.

The G58R substitution lies within a conserved hydrophobic α-helix of C10 and contributes to a GXXXGXXXG motif that often mediates protein-protein interactions (Huang et al., 2018; Russ and Engelman, 2000). The G58R substitution, which introduces a positive charge into the predicted hydrophobic glycine cleft of the GXXXGXXXG helix, is likely to disrupt the α-helix and/or its interactions. To test this idea further we deleted the α-helix of C10 (ΔαH). Similar to C10 G58R, C10 (ΔαH) induced the fragmentation of mitochondria in an OMA1-dependent manner (Figure 3D and E). These findings are consistent with the notion that the C10 G58R mutation disrupts the α-helix of C10 to induce OMA1/OPA1 dependent mitochondrial fragmentation.

### Intense C2/C10 foci accumulate on blocking L-OPA1 cleavage by OMA1

Unexpectedly, we observed that wildtype C10-Flag formed foci within OMA1 KO cells, rather than the typical smooth appearance throughout the mitochondrial network of overexpressed C10 protein by light microscopy (Figure 3A). To investigate whether the pattern of endogenous C2 and C10 distribution also changes in the absence of OMA1, we used antisera specific for these proteins in OMA1 KO HeLa cells (Figure 4A – C and Supplemental Figure 3A - B). We previously reported that endogenous C2 and C10 form foci in the intermembrane space that extend into the mitochondrial cristae in WT HeLa cells (Huang et al., 2018). Strikingly, in the absence of OMA1, C2 and C10 foci exhibited substantially increased in intensity (Figure 4A – C and Supplemental Figure 3A - B). As the pattern was similar for both C2 and C10, we focused on C2 in subsequent analyses. High intensity foci were present in the majority of OMA1 KO cells and absent in OMA1 KO cells re-expressing WT but not protease-dead OMA1 (Figure 4A and B). Thus, the pattern of C2 and C10 distribution is greatly altered in the absence of OMA1 with intensification of C2/C10 foci in the mitochondrial network.

**Figure 4:**
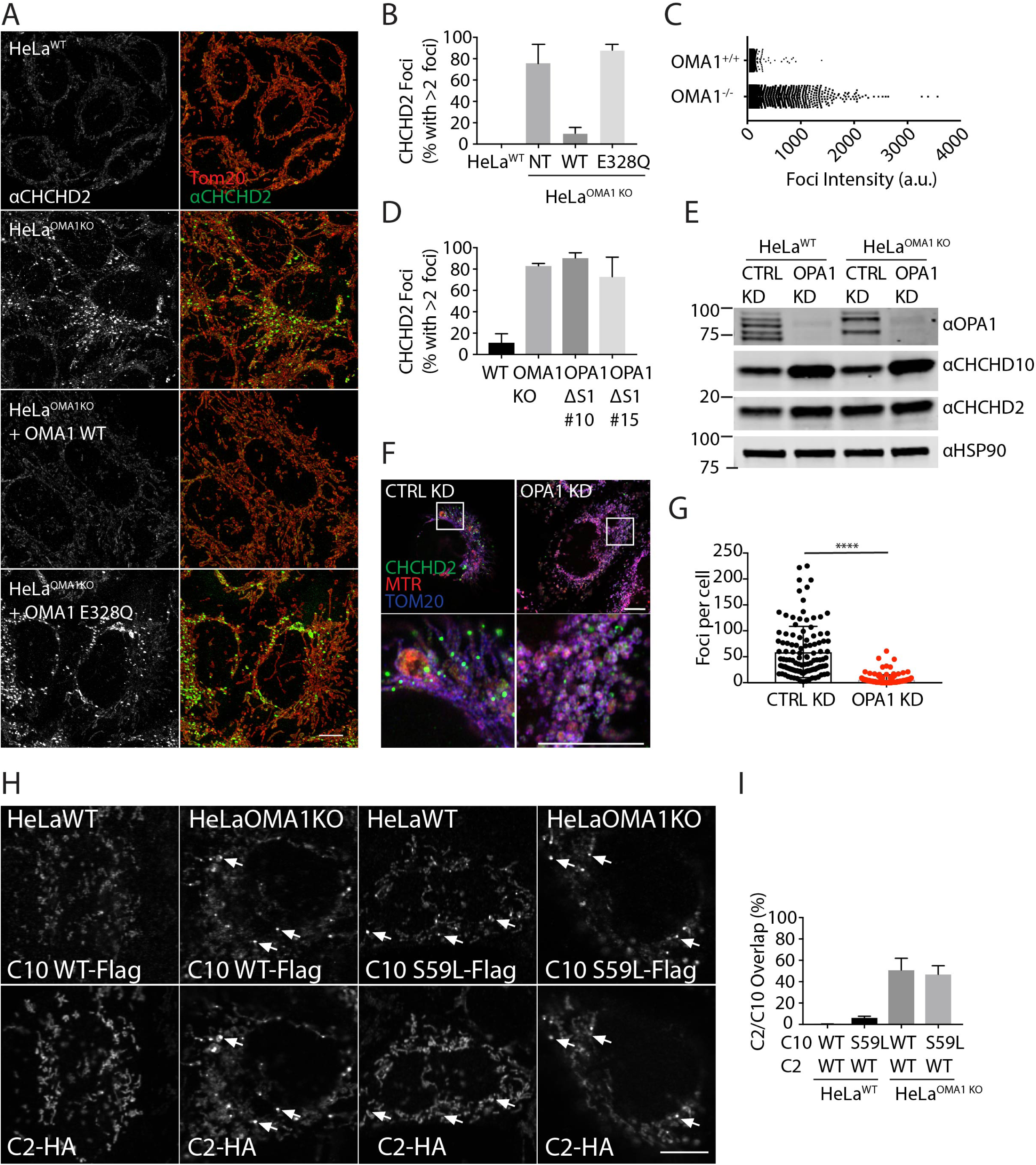
Altered C2/C10 localization and processing in the absence of OMA1. (A) Airyscan confocal images from of WT and OMA1 KO HeLa Cells immunostained for Tom20 (red) and C2 (green). Where indicated OMA1 KO HeLa cells stably expressed OMA1 WT or the protease-dead OMA1 E328Q mutation. Scale bar = 20 μm. (B) Quantification of cells with greater than 2 intense foci in (A). Greater than 150 cells were scored in 3 biological replicates. (C) Graph depicts intensities of individual foci from representative WT and OMA1 KO HeLa cells. (D) Quantification of cells with greater than 2 intense C2 foci in WT or two independent OPA1Δs1 HeLa cell lines (#10 and #15). >150 cells were scored in three biological replications for each condition. (E) Immunoblot of WT and OMA1 KO HeLa cells treated with control or OPA1 siRNA as in (A). HSP90 served as the loading control. Immunoblot is representative of 3 biological replicates. (F) Representative Airyscan confocal images of OMA1 KO HeLa cells treated with control or OPA1 siRNA and stained for C2 (green), Mitotracker Red (MTR) (red), and Tom20 (blue). (G) Quantification of intense C2 foci in OMA1 KO Hela cells following treatment with either control or OPA1 siRNA. >150 cells were scored in three biological replications for each condition. “****” indicates “p-value < 0.0001”. (H) Representative confocal images of WT or OMA1 KO HeLa cells transient co-transfected with CHCHD2-HA and either CHCHD10(WT)-Flag or CHCHD10(S59L)- Flag, immunostained for Flag and HA. (I) Quantification of (H) for CHCHD10 foci that overlap with CHCHD2 foci. > 20 cells in each condition were analyzed. Scale bars in all images = 10 μm. All confocal images were acquired as a z-stack and represented as a maximum projection image.

Given OPA1 processing is altered in OMA1 KO cells, we next investigated the dependence of high intensity C2 foci on OPA1 and its processing. To assess whether C2 foci formation depends on OPA1 processing by OMA1, we used genomic editing to delete the canonical OMA1 cleavage site within the endogenous OPA1 locus (HeLa^OPA1Δs1^). This blocked formation of the canonical OMA1 cleaved products *c* and *e* — although residual degradation of OPA1 was observed following mitochondrial depolarization by CCCP, possibly due to limited cleavage by OMA1 at a non-canonical site (Supplemental Figure 3D). Importantly, C2 foci were observed also in HeLa^OPA1Δs1^ cells, indicating that partially blocking cleavage of OPA1 by OMA1 leads to formation of high intensity C2 foci (Figure 4D). To further assess the dependence of C2/C10 formation on OPA1, we depleted OPA1 using siRNA (Figure 4E). Interestingly, C10 levels increased following OPA1 knockdown in HeLa cells (302 ± 75% in HeLa^WT^ and 325 ± 85% in HeLa^OMA1KO^, p = 0.0018). Despite an increase in total C2 protein levels, C2 foci were found to be significantly decreased, suggesting that their formation is a consequence of altered OPA1 isoform balance and not a separate function of OMA1 (Figure 4F and G). Together these findings are consistent with the notion that C2 foci accumulate due to failure of OPA1 processing by OMA1.

The C2/C10 foci resembled those that we previously observed to form spontaneously with transient expression of the C10 mutant S59L (Huang et al., 2018). To further explore the relationship between these foci, we co-expressed C2(WT)-HA with C10(WT)-Flag or C10(S59L)-Flag. Only C10(S59L)-Flag formed foci in wildtype HeLa cells; but, notably, failed to recruit C2(WT)-HA into the foci (Figure 4H and I). By contrast, both C10(S59L)-Flag and C10(WT)-Flag formed foci in similar numbers in HeLa OMA1 KO cells. The majority of these foci also contained C2-HA. Although not conclusive, we hypothesize these intense foci may represent stalled intermediates that can form either from failure of C10(S59L) to disperse, due to increased hydrophobicity in the N-terminal αH, or from blocked OPA1 cleavage in the absence of OMA1.

### A pool of C2/C10 is degraded following activation of OMA1 by mitochondrial stressors

We and others previously demonstrated that C2 and C10 exhibit short half-lives in cultured cells (Burstein et al., 2018; Huang et al., 2018). To test whether OMA1 might be the protease responsible for basal C2/C10 degradation, and whether this might account for C2/C10 foci formation, we first assessed C2/C10 protein levels after inhibition of protein translation with cycloheximide (Figure 5A). CHX treatment times were chosen based on our prior work (Huang et al., 2018). C2/C10 half-lives were similar in wildtype and OMA1 KO HeLa cells, demonstrating that OMA1 is not required for basal turnover of C2/C10. C2/C10 half-lives were similarly unaffected by loss of YME1 or PARL, two inner membrane proteases in complex with OMA1 and the anchoring protein SLP-2 (Supplemental Figure 4A) (Wai et al., 2016).

**Figure 5:**
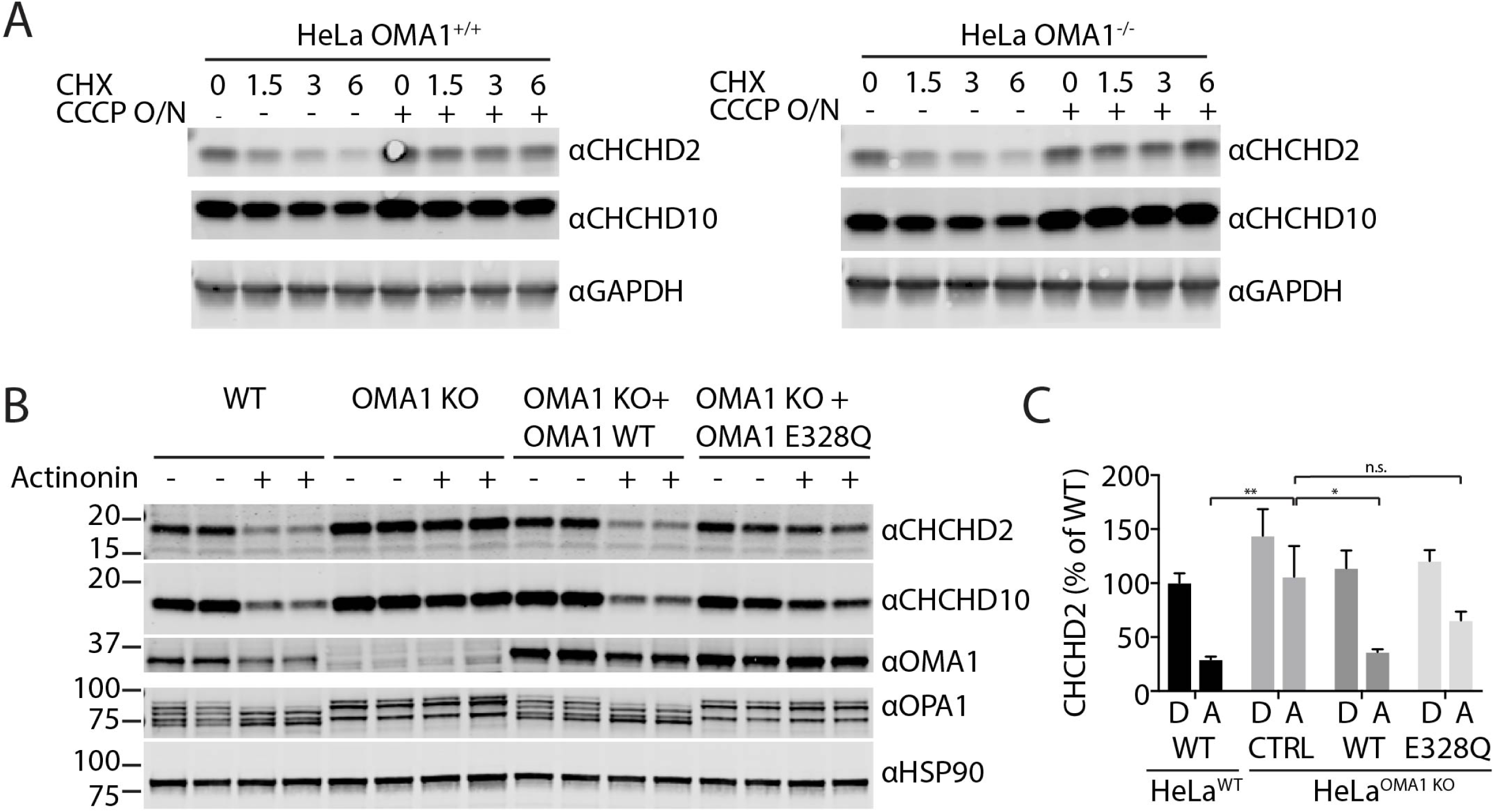
C2/10 are degraded by OMA1 activated by mitochondrial stress. (A) Immunoblot of OMA1 KO and WT HeLa cells treated with DMSO or 10 μM CCCP overnight (O/N) and then treated with 100 μM cycloheximide (CHX) for the indicated number of hours. GAPDH or Tubulin served as controls where indicated. (B) Immunoblot of WT and OMA1 KO HeLa cell lines, which stably express OMA1 WT or OMA1 E328Q mutation where indicated, treated with either vehicle or 125 µM actinonin for 3 hours. HSP90 served as the loading control. (C) Quantification of the C2 levels from (B). N = 3 biological replicates per sample. The levels of C2 of each cell type following DMSO (D) or actinonin (A) treatment are normalized to those from WT HeLa with DMSO treatment. Three biological replicates were quantified.

We next tested cells subject to mitochondrial stressors known to activate OMA1. Actinonin treatment for 3 hrs has been previously shown to accumulate aberrant mtDNA-encoded polypeptides in the inner membrane and potently activate OMA1 prior to collapse of the mitochondrial membrane potential (Richter et al., 2015). Unexpectedly, we found that C2/C10 levels are reduced following 3 hr treatment with actinonin (Figure 5B and C and Supplemental Figure 4B). This reduction was dependent on OMA1, as it was partially blocked in OMA1 KO cells. Consistently, degradation was restored in OMA1 KO cells stably re-expressing WT but not catalytically inactive OMA1. Together these findings suggest that OMA1 is dispensable for basal turnover of C2/C10, but, when activated by inner membrane proteostatic stress, degrades C2/C10.

OMA1 is also potently activated by mitochondrial uncoupling with drugs such as CCCP (Ehses et al., 2009; Head et al., 2009). As we reported previously and in contrast to treatment with actinonin, C2/C10 levels increased in response to loss of membrane potential (Huang et al., 2018). This increase in C2/C10 abundance with uncoupling was independent of OMA1, as it occurred in both WT and OMA1 KO HeLa cells (Figure 5A). Co-treatment with CCCP and actinonin resulted in accumulation of C2/C10 similar to treatment with CCCP alone (Supplemental Figure 4C). Thus, although C2/C10 is degraded following OMA1 activation in the presence of a mitochondrial membrane potential, the pool of C2/C10 that accumulates with loss of membrane potential is protected from OMA1 and other proteases.

Together these findings suggest that OMA1 degrades C2/C10 upon activation but is not responsible for their basal turnover. This also implies that the formation of C2/C10 intense foci in the absence of OMA1 is not a direct consequence of failed C2/C10 degradation by OMA1, but rather reflects loss of OMA1 activity against another substrate (e.g., L-OPA1).

In summary, these findings suggest that the functional interaction between C2/10 and OPA1/OMA1 is bidirectional. C2/10 loss or C10 mutation leads to OMA1 activation and OPA1 processing. Conversely, OMA1 loss blocks the stress-induced degradation of C2/10; and C2/10 collect in intense foci in OMA1-deficient mitochondria.

### Pathogenic C10 mutation activates OPA1 cleavage by OMA1 in vivo

Having identified a functional interaction between pathogenic C10 mutants and OMA1 activation in cell culture, we next asked whether mutant C10 activates OMA1 *in vivo*.

To model the pathogenic S59L mutation in human C10, we generated a transgenic mouse with the equivalent substitution in mouse C10 (S55L in mouse; but, for clarity, numbering corresponding to human C10 is used throughout) (Figure 6A). Consistent with two recent reports on independent lines (Genin et al., 2019; Anderson et al., 2019), we found that C10^S59L/+^ mice developed cardiomyopathy by early adulthood (17 – 42 weeks). % ejection fraction (%EF) in C10^S59L/+^ animals was significantly lower than WT littermates by echocardiography (39.6% vs 59.7%, p = 0.003) (Figure 6B – D and Table S2). Similarly, peak left ventricular outflow tract (LVOT) and pulmonary artery (PA) velocities were significantly reduced, indicating, respectively, left and right ventricular dysfunction (624.1 vs. 1204 mm/s, p < 0.0001; and 424.5 vs. 686.5 mm/s, p < 0.0001, respectively). Additionally, the anterior and posterior left ventricular walls were significantly thickened in diastole by echocardiography (Supplemental Figure 5 and Table S2). Vacuole formation in cardiac tissue and fibrosis were apparent on histology with H&E and Masson trichrome staining, respectively (Figure 6E). C10 distribution was also found to be greatly altered in cardiac tissue from C10^S59L/+^ knock-in mice, forming foci resembling those observed in cell culture (Figure 6F) (Huang et al., 2018). Together these findings demonstrated development of hypertrophic cardiomyopathy in C10^S59L/+^ knock-in mice by early adulthood.

**Figure 6:**
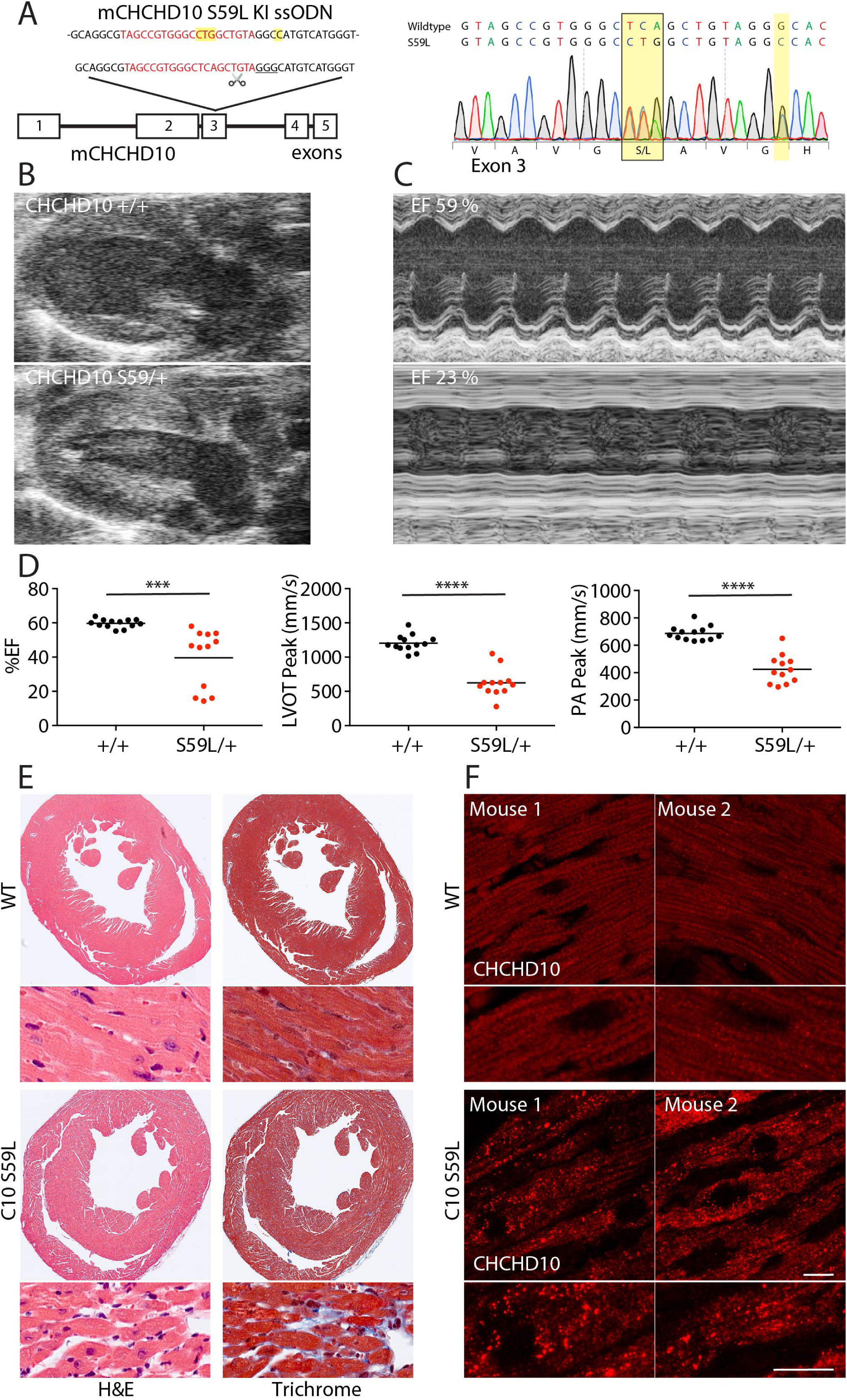
C10 ^S59L/+^ KI mice develop cardiomyopathy. (A) C10^S59L/+^ KI mice were generated using CRIPSR/Cas9 genomic editing. The targeted CRISPR cut site (scissors), guide RNA sequence (red), and PAM sequence (underlined) are indicated. The donor ssODN used for repair is shown above. Sanger sequencing demonstrates the TCA -> CTG codon change, as well as a silent GGG -> GGC mutation at G58 introduced to disrupt the PAM sequence. (B) Representative echocardiogram in long axis view from WT and C10^S59L/+^ mice demonstrating thickening of the ventricular wall in C10^S59L/+^ mice. (C) Motion (M)-Mode view from the short axis of echocardiogram, demonstrating decreased left ventricular contractility of the C10^S59L/+^ heart. (D) Graph representing % ejection fraction (%EF), left ventricular outflow tract velocities (LVOT), and pulmonary artery velocities (PA) of litter matched WT (N = 13) and C10^S59L/+^ (N = 12) mice between the ages of 17 - 42 weeks. Velocities were measured by pulsed-wave Doppler echocardiography. “***” denotes p-value < 0.001 and “****” indicates p-value < 0.0001. (E) Representative images of H&E stain (left) and Masson trichrome stain (right) from adult WT and C10^S59L/+^ mice. (F) Airy scan confocal images of C10 immunostaining of heart slices from WT and C10^S59L/+^ mice. Scale bar = 10 μm. All confocal images were acquired as a z-stack and represented as a maximum projection image.

Examining cardiac tissue lysates from C10^S59L/+^ mice, we observed a substantial elevation in C2 and C10, consistent with a recent report *in vivo* and our prior observation that C2 and C10 increase in response to mitochondrial distress in cell culture (Figure 7A) (Huang et al., 2018; Anderson et al., 2019). Interestingly, C2 and C10 protein levels decreased slightly with aging in wildtype mice. MTHFD2 and MTHFD1L, which are sensitive markers of the mitochondrial integrated stress response (mt-ISR) in muscle (Kühl et al., 2017; Khan et al., 2017), also progressively increased from younger (9 - 13 weeks) to older (22 – 37 weeks) C10^S59L/+^ animals (Figure 7A). Consistent with activation of the mt-ISR in C10^S59L/+^ mice, expression of transcription factors reported to mediate the response in cardiac tissue, ATF4, ATF5, CHOP, and Myc (Kühl et al., 2017), were significantly elevated relative to younger wildtype mice (Figure 7B). Altogether, in the younger C10^S59L/+^ vs. younger WT comparison, altered expression reached gene-level significance for 15 of 22 pre-specified mt-ISR-associated genes, recently identified to be differentially regulated genes (DEGs) in the heart of an independent C10 ^S59L/+^ mouse line (Supplemental Figure 6 and Table S3) (Anderson et al., 2019).

**Figure 7:**
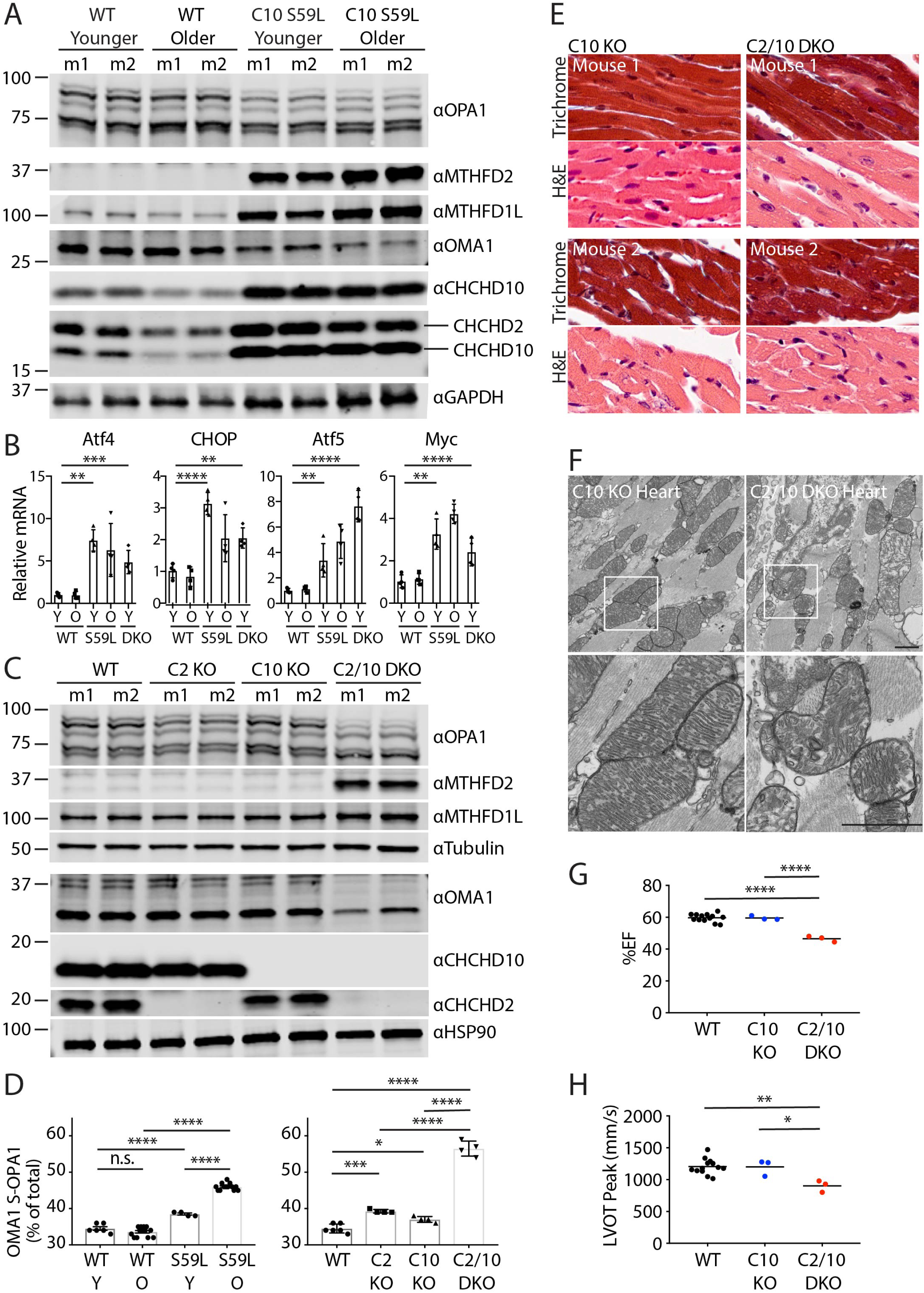
OMA1 activation co-incident with cardiomyopathy and mt-ISR phenocopied in C2/C10 DKO and C10^S59L/+^ KI mice. (A) Representative immunoblot of heart lysates from younger (9 - 13 weeks) and older (36 – 37 weeks) WT and C10^S59L/+^ KI mice. Lysates from two different mice (m1 and m2) are shown for each condition. (B) Relative transcript levels of Atf4, CHOP, Atf5, and Myc in heart extracts of younger WT, C10^S59L/+^ KI, and C2/C10 DKO mice (9 - 13 weeks), and older WT and C10^S59L/+^ KI (22 - 37 weeks) mice measured by microarray and normalized to average of younger WT mice. N = 4 for each group. (C) Representative immunoblot of heart lysates from WT, C2 KO, C10 KO, and C2/C10 DKO mice (age 9 - 13 weeks). Lysates from two different mice (m1 and m2) are shown for each condition. (D) Quantification of OMA1-cleaved S-OPA1 relative to total OPA1 from heart lysates as in (A and C). Note: WT comparison group is same as in the right and left graphs. (E) Representative images of H&E stain and Masson trichrome stain from sibling-matched C10 KO and C2/C10 DKO mice (9 - 13 weeks). (F) Representative TEM images of C10 KO and C2/C10 DKO hearts. (G and H) Graph representing echography measurements of % ejection fraction (%EF) (G) and left ventricular outflow tract velocities (LVOT) (H) for sibling matched C10 KO and C2/C10 DKO mice and WT mice in (Fig. 5D). C10 KO and C2/C10 DKO mice were 22 – 28 weeks old at the time of echocardiography. In all graphs, “n.s.” indicates “not significant,” “*” corresponds to p-value < 0.05, “**” represents p-value < 0.01, “***” denotes p-value < 0.001 and “****” indicates p < 0.0001.

We next assessed OPA1 processing in C10^S59L/+^ mice. Notably, in the heart, L-OPA1 (isoforms *a* and *b*) was excessively processed to OMA1 cleavage products (isoforms *c* and *e*) in younger C10^S59L/+^ mice and to an even greater extent in older C10^S59L/+^ mice (Figure 7A and D). OMA1 levels were likewise decreased in C10^S59L/+^ hearts, as expected given OMA1 is degraded shortly after its activation (Head et al., 2009). Sampling lysates from the atria and ventricles, separately, demonstrated that the myocardium is similarly affected throughout (Supplemental Figure 7A). OMA1 activation was absent in the liver (Supplemental Figure 7B). C10 and MTHFD2 protein levels were also normal in liver; thus, OMA1 activation correlates with other signs of mitochondrial distress *in vivo*. Notably, OMA1 was not activated in either C10^S59L/+^ or C10^S59L/S59L^ primary fibroblast lines, in contrast to C2/C10 DKO cell lines, suggesting that the pathogenic effect of mutant C10 may require its high expression in the heart (Supplemental Figure 7C). Together these findings demonstrate that the pathogenic C10 mutation activates OPA1 cleavage by OMA1 *in vivo*, possibly following its accumulation with C2 in response to an inciting mitochondrial stress.

### C2/C10 double knockout activates OPA1 cleavage by OMA1 and mt-ISR in vivo

Guided by parallel results in DKO fibroblasts and C10^S59L/+^ mutant hearts, we assayed OPA1 processing in lysates from younger C2/C10 single and double knockouts mice (9 – 13 weeks). Whereas processing of OPA1 to OMA1-cleaved forms was mildly increased in hearts from C2/C10 single knockout mice, loss of both C2 and C10 had a synergistic effect on L-OPA1 cleavage (Figure 7C and D; Figure 1G). OMA1 levels were likewise decreased in DKO hearts, as expected given OMA1 is degraded shortly after its activation. OMA1 cleavage of L-OPA1 was also apparent in liver, brain, and skeletal muscle at this age, albeit to a lesser extent (Supplemental Figure 8A and B). Comparing OMA1 activation in the younger C10^S59L^ model to the younger knockout models, showed OMA1 activation in C10^S59L^ to be intermediate between C10 single KO and C2/C10 DKO (Figure 7D). Together these findings demonstrated that OMA1 is activated by C2 and C10 loss *in vivo*, phenocopying OMA1 activation by C10^S59L/+^.

It was recently reported that C10 KO does not activate the mt-ISR in the mouse heart, in contrast to C10^S59L/+^ KI. Consistently, we did not see an increase in MTHFD2 or MTHFD1L in hearts of either C10 or C2 single KO mice (Figure 7C). By contrast, double knockout of C2 and C10 led to a substantial elevation in MTHFD2 and a more modest elevation in MTFHD1L protein levels. MTHFD2 was similarly activated in skeletal muscle and brain but not apparent in liver (Supplemental Figure 8A and B). Consistently, expression of transcription factors associated with the mt-ISR (namely, ATF4, ATF5, and Myc) were significantly elevated in younger DKO mice compared to younger WT mice with CHOP reaching nominal significance (Figure 7B and Table S3). Altogether, 6/22 genes in a pre-specified set of mt-ISR associated genes previously identified as DEGs in the C10^S59L/+^ KI model reached gene-level significance in the overall DKO vs. WT comparison (Figure 7B and Supplemental Figure 6). With the exception of ATF5 expression, generally the mt-ISR was less robust in C2/C10 DKO mice compared to C10^S59L/+^ KI mice examined at a similar age. Comparison of total DEGs between DKO vs. WT and C10^S59L/+^ vs. WT also underscored the similarity in overall transcriptional response and the greater severity in the younger C10 ^S59L/+^ mice compared to younger DKO mice. The C10 ^S59L/+^ vs. WT comparison revealed 1,386 DEGs, whereas the C2/C10 DKO vs. WT comparison produced 548 DEGs, 290 (52.9%) of which were shared. Thus, a similar transcriptional response occurs in both C2/C10 DKO and C10^S59L/+^ KI hearts but involves a more extensive set of genes in C10^S59L/+^ KI hearts.

Given early biochemical changes in the heart, we next assessed our younger DKO mice for early signs of cardiomyopathy. Although fibrosis was not apparent by 13 weeks, numerous vacuoles similar to those seen in C10^S59L/+^ KI were observed in tissues from DKO mice with both H&E and Masson Trichrome staining (Figure 7E). These changes were not apparent in single C10 KO litter mates. Additionally, consistent with TEM findings in C10 KO and C2/10 DKO primary fibroblasts and L-OPA1 cleavage patterns in the heart, TEM cristae structure was abnormal in the majority of mitochondria from a C2/10 DKO heart but normal in most mitochondria from a litter-matched C10 KO heart (96/116 [82.8%] abnormal vs. 11/143 [7.6%] abnormal) (Figure 7F). To assess for cardiomyopathy, we evaluated our oldest cohorts of C2/C10 DKO and C10 KO littermates by echocardiography. By 22 - 29 weeks, significantly decreased % ejection fraction was apparent in DKO mice when compared to both their C10 KO littermates and unrelated WT mice of the same strain (46.5% vs. 59.53% and 46.5% vs. 59.68%, p-value < 0.0001 for both comparisons) (Figure 7G and Table S2). Similarly, LVOT peak velocity and PA were decreased in DKO mice compared to their C10 KO littermates and unrelated WT mice (902.7 vs 1199 cm/s, p = 0.0130; and 540.7 vs 704.7 cm/s, p = 0.0014), consistent with left and right ventricular dysfunction, respectively (Figure 7H, Supplemental Figure 8C, and Table S2). Cardiac dysfunction in C2/C10 DKO mice appeared to be less severe than in unrelated C10^S59L/+^ KI mice, with the caveat that relatively few similarly aged mice were available for comparison. Thus, loss of C2 and C10 causes early cardiomyopathy, partially phenocopying C10^S59L/+^ KI mice.

Thus, both loss of C2/C10 and mutation in C10 cause OMA1 activation in the mitochondrial intermembrane space (Figure 8, top). Activated OMA1 cleaves the cristae shaping and mitochondrial fusion protein L-OPA1 into its short form, destabilizing cristae structure. As OPA1 is known to be necessary for mtDNA stability, we speculate that OPA1 cleavage by OMA1 may be important for mtDNA instability and mt-ISR reported in patients and the C10 ^S59L/+^ mouse (Bannwarth et al., 2014; Genin et al., 2019).

**Figure 8.**
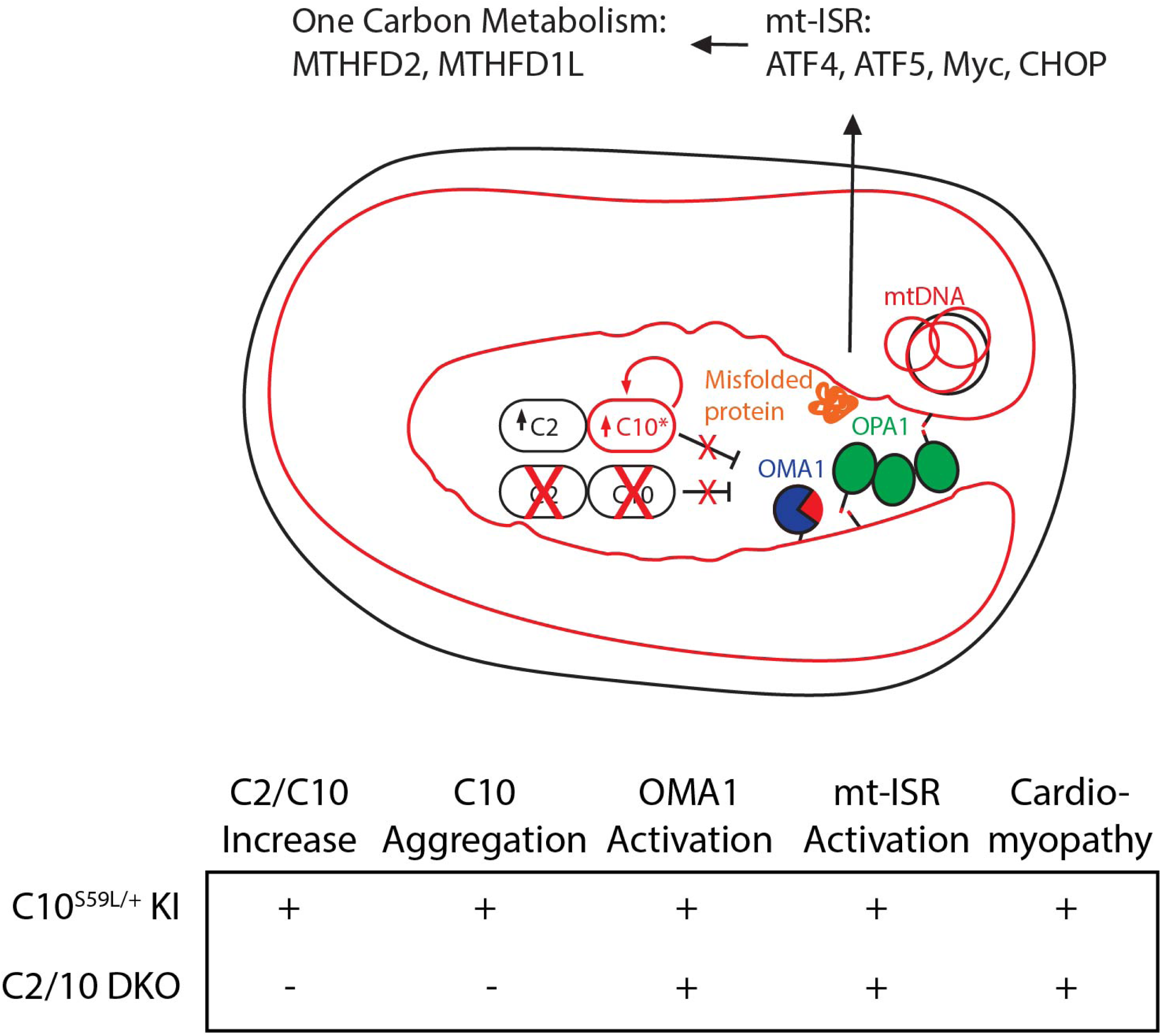
Model of OMA1 activation due to loss of C2/C10 and mutant C10. (Top) Model representing OPA1 processing by OMA1 in the setting of C2/C10 loss and C10 mutation in complex WT wildtype C2. Activated OMA1 cleaves OPA1. We speculate that OPA1 cleavage may impact mtDNA stability and mitochondrial integrated stress response (mt-ISR) observed in C10^S59L/+^ mice. A transcriptional target of the mt-ISR are mitochondrial proteins involved in one-carbon metabolism, including MTHFD1L and MTHFD2. (Bottom) Table illustrates findings shared and not shared by the C10^S59L/+^ and C2/10 DKO mice. Notable, OMA1 activation, mt-ISR, and cardiomyopathy are common to the models. However, C2 and C10 protein increase and aggregation are unique to C10^S59L/+^ mice. This suggests that C10^S59L/+^ causes OMA1 activation, mt-ISR, and cardiomyopathy by impinging on the normal function of C2/C10; it argues against misfolded C10 protein having only a toxic effect of mitochondria independent of its normal function.

Comparison of the C10 ^S59L/+^ and C2/10 DKO mice also challenges at least the simplest formulation of the C10 toxic aggregation model, which supposes that C2/C10 toxic aggregates directly activate mt-ISR and lead to tissue degeneration independently on their normal function (Figure 8, bottom) (Anderson et al., 2019). That no C2/C10 is present to aggregate in the DKO mouse and yet it phenocopies (at least in part) the mt-ISR and cardiomyopathy of the C10^S59L/+^ mouse suggests that mutant C10 pathogenesis is closely related to the normal function of C2/C10.

## Discussion

In this study, we identified a functional interaction between C2/C10 and stress-induced processing of L-OPA1 by OMA1 that is shared by C2/C10 loss of function and mutant C10 gain-of-function.

Examining primary cells from the first C2/C10 DKO mouse we found cristae to be decreased and occasionally circular and detached from the boundary membrane. Morphology in single KO cells was normal, consistent with functional redundancy between the paralogs in mammals. These findings are consistent with swirled cristae recently reported in *Drosophila* muscle lacking the predominate C2/C10 ortholog in that tissue, CG5010 (Meng et al., 2017). They are also reminiscent of cristae abnormalities observed in fibroblasts and skeletal muscle from patients with C10 mutations (S59L and G58R, respectively) (Ajroud-Driss et al., 2015; Bannwarth et al., 2014).

As an explanation for cristae abnormalities, C10 (and more recently C2) have been proposed to be structural subunits of the MICOS complex, which supports cristae integrity by bending the inner membrane at the cristae junction. However, a number of observations have challenged this view. First, the bulk of C2 and C10 in the cell migrates in a ∼200 kDa complex and not with the MICOS complex by BN-PAGE and sucrose gradient (Straub et al., 2018; Burstein et al., 2018). Second, unlike MICOS subunit MIC19, endogenous C2 and C10 are not restrained at the cristae junction but extend into the cristae by immuno-EM (Huang et al., 2018). Third, unlike MICOS subunits, MIC19, MIC26, and MIC27, C2 and C10 are stable upon knockdown of the core MICOS subunit, MIC60. Fourth, whereas MIC19 co-immunoprecipitates MIC26 and MIC27, C2 fails to do so under the same conditions (Huang et al., 2018). In this report, we extended these previous findings by establishing that that the MICOS complex is stable in the absence of C2 and C10. Additionally, we found that fibroblasts from C2/10 DKO and C10 KO siblings have similar expression of MIC19, MIC26, MIC27, and MIC60, despite the former but not the later having disrupted cristae by TEM.

As an alternative explanation for cristae abnormalities in the absence of C2/C10 we identified L-OPA1 cleavage by activated OMA1. This was observed in HEK293 cells, primary fibroblasts, and heart and skeletal muscle *in vivo*. We extended these findings to mutant C10, identifying OMA1 processing of OPA1 as a key event in affected organs, such as the heart. Thus, we identify L-OPA1 cleavage by OMA1 as a novel mechanism for cristae abnormalities resulting from C10 mutation or C2/C10 loss.

Loss of L-OPA with mutant C10 S59L may also explain other aspects of the phenotype. Affected muscle in patients and a mouse model with C10 S59L mutation harbored multiple mtDNA deletions (Bannwarth et al., 2014; Genin et al., 2019), which have also been detected in muscle from patients with OPA1 mutation (Hudson et al., 2008). Future studies modeling C10 mutations on an OMA1 KO background in mice will likely help assess the impact of L-OPA1 on cristae structure and mtDNA instability.

The trigger for OMA1 activation in the absence of C2 and C10 is not clear at present. We failed to detect direct physical interactions between C2 and OPA1 or OMA1 stable enough for immunocapture, and at least the predominate pool of C2/C10 appears in complexes that are of distinct in size from the predominate OPA1- and OMA1-containing complexes by BN-PAGE (Frezza et al., 2006; Wai et al., 2016; Straub et al., 2018). Thus, activation of OMA1 with C2/C10 loss is more likely indirect, perhaps by causing a specific mitochondrial stress.

The two best characterized stressors activating OMA1 are bioenergetic collapse and proteostatic stress following overload of quality control (QC) proteases (Head et al., 2009; Ehses et al., 2009). OMA1, for example, is activated following loss of AAA proteases YME1, AFG3L2, and SPG7, as well as scaffolding proteins, such as prohibitins and SLP-2, that organize YME1 and AFG3L2/SPG7, respectively, into microdomains in the inner membrane (Merkwirth et al., 2008; Ehses et al., 2009; Anand et al., 2014; Wai et al., 2016; Magri et al., 2018). Similarly, overload of QC proteases with aberrant mtDNA translation products in the setting of actininon treatment potently activates OMA1 prior to membrane potential collapse (Richter et al., 2015).

Under growth conditions in which OMA1 was activated in C2/C10 DKO cells, oxygen consumption and membrane potential were minimally affected, suggesting that OMA1 activation in these cells is not primarily driven by bioenergetic stress. An alternative possibility that loss of C2/C10 triggers OMA1 activation by increasing proteostatic stress is intriguing and has some circumstantial evidence in support (although direct evidence is lacking). Previous studies have identified interactions between either C2 or C10 and the proteases YME1 and PARL, which are components of the intermembrane space facing SPY complex (Floyd et al., 2016; Wai et al., 2016). We were able to replicate an interaction between C2 and YME1 in this study (although interactions with PARL and SLP-2 were not observed under our study conditions). These findings are at least consistent with a transient interaction between C2/C10 and components of this complex of proteases. Additionally, we observed that in the absence of OMA1, C2/C10 form intense foci in unusual mitochondria dilatations containing a complex arrangement of inner membrane. We speculate that these mitochondrial dilatations result from failure of OMA1 to trigger L-OPA1 cleavage in response to local proteostatic stress. Consistently, similar mitochondrial dilatations in OMA1 KO cells have been reported to accumulate with actinonin treatment, following an increase in inner membrane proteostatic burden (Richter et al., 2015). We hypothesize that these intense C2/C10- containing foci located in cristae represent stalled intermediates accumulating in areas of local proteostatic stress -- although alternative explanations, such as their association with a particular inner membrane structure or fusion intermediate are also plausible. Identification of other proteins present in these foci will likely clarify their nature. Finally, as we and others have observed previously, C2/C10 have short half-lives and are dynamically regulated with a dramatic increase in IMS expression following loss of the inner membrane potential (Burstein et al., 2018; Huang et al., 2018). Whereas this dynamic regulation seems incompatible with a primarily structural role, it would be advantageous for proteins responding to proteostatic stress.

Early changes in C2/C10 DKO mice suggest that they partially phenocopy C10^S59L/+^ mice, with development of cardiomyopathy co-incident with activation of OMA1 and the mt-ISR in the heart. Although a haploinsuffiency model is ruled out by the normal phenotype in the C10 single KO mouse, this implies that the physiologic function of C2/C10 and mutant C10 pathogenesis may be related, nevertheless. In the most straightforward model, mutant C10 may have a dominant negative effect on the C2/C10 complex in which it assembles, inhibiting C2/C10 function. Alternatively, or in addition, C10 S59L may cause disease by a toxic gain-function mechanism that is related to its normal function, e.g., through dysregulation of a physiologic interaction. Despite their similarities there are some differences in the severity of changes observed in the C10^S59L/+^ KI and C2/C10 DKO mice, with greater OMA1 activation in the C2/C10 DKO mouse and greater mt-ISR activation and more severe cardiomyopathy in the C10^S59L/+^ KI mouse. This may be most consistent with a mixed model in which accumulated C10^S59L/+^ has a dominant negative effect on the C2/C10 complex thereby triggering OMA1 activation and additionally has a toxic effect on mitochondria. In this model, both dominant negative and toxic gain-of-function effects would contribute to pathogenesis.

Identifying OMA1 activation as a key event in C10 mutant pathogenesis, also nominates OMA1 as a potential therapeutic target for C2/C10-related disorders. Although OMA1 is required for OPA1 processing in response to mitochondrial stress, it is not essential for mammalian life (Quirós et al., 2012). The OMA1 KO mouse is viable and loss of function mutations occur in human population at close to the rate predicted by chance, including two “knockout” adult individuals with homozygous frameshift mutations (observed SNV/expected SNV = 0.78 (0.55 – 1.13); gnomAD v.2.1.1 database) (Karczewski et al., 2019). Indeed, in the setting of a high proteostatic load, such as that resulting from loss of YME1 or the scaffolding protein prohibitin-2, its excessive activation may be detrimental, leading to heart failure and neurodegeneration, respectively (Wai et al., 2015; Korwitz et al., 2016). Future studies of mutant C10 (and C2) on an OMA1 knockout background will be important for evaluating OMA1 as a therapeutic target, as well as determining which aspects of the mutant phenotype (e.g., cardiomyopathy, mt-ISR activation, multiple mtDNA deletion, cytosolic TDP-43 accumulation) are dependent on L-OPA1 processing by OMA1.

## Methods

### Generation of transgenic mice

C2^-/-^, C10^-/-^, and C10^S59L/+^ transgenic lines were produced using CRISPR/Cas9 endonuclease-mediated genome editing on the C57Bl6J background. CRISPR guide RNAs targeted exon 2 and 4 in mC10 and exon 1 and 4 in mC2 to generate breaks in DNA. Animals were screened for large deletions lacking the aforementioned exons by Sanger sequencing. C10 KI mice were generated with a guide RNA targeting near the S55 position (equivalent of S59 in human C10) in mC10 co-injected with Cas9 and an ssODN containing a TCA -> CTG substitution at the S55L codon in the C57Bl6J background. All animal studies were approved by the Animal Care Use Committee at the NINDS intramural program. Both genders were used in all studies with the exception of echocardiography of the C10 KO and C10 DKO for which only female animals were used given availability of appropriately aged animals in the colony for this testing.

### Cell Culture

WT, C2, C10 and DKO fibroblast cells were generated from 1 day old newborn pups. Pups were removed from litter and placed in DMEM containing 1% L-glutamine, 15% FBS, MEM-NEAA, 1% sodium pyruvate, 1X pen/strep, 1X gentamicin, and 1X amphotericin B. Sterile scalpels were used to cut the skin from the animals into thin slices around 1 - 2mm squares. The tissues were transferred to a 1% gelatin-coated plate containing the dissection media and covered with sterile cover slips. High glucose DMEM with sodium pyruvate and supplemented with 10% FBS and pen/strep was used for all other tissue culture and for primary fibroblasts after passage 1.

OPA1 null Mouse Embryonic Fibroblasts were obtained from ATCC (ATCC CRL-2995) and transformed MEF WT cells with SV40 were obtained as a kind gift from the laboratory of David Chan (Caltech). Generation of the HEK293 C2/C10 DKO cell line was described previously (Huang et al., 2018). HeLa^PARL^ KO, HeLa^OMA1 KO^, HeLa^OMA1 KO^ stably expressing OMA1 WT or OMA1 E328Q and matched HeLa^WT^ cell lines were a kind from Richard Youle (NIH) and have been described previously (Sekine et al., 2019). HeLa^YME1 KO^ and matched HeLa^WT^ cell lines were a kind gift from Thomas Langer (Cologne) and have been described previously (Hartmann et al., 2016).

HEK293 OMA1 KO and HEK293 C2/C10/OMA1 TKO lines were produced by CRISPR/Cas9, using the pSpCas9(BB)-2A-GFP (PX458) plasmid with the guide sequences 5’ CACCGAGTGCAATCGAGACGTCCCG 3’ and 5’ CACCGAGATCGCACACGCAGTCCTG 3’ targeting exons 4 and 5, respectively. Knockout was verified by immunoblotting for OMA1 and characteristic change in OPA1 processing. The HeLa^OPA1Δs1^ was produced using the pSpCas9(BB)-2A-GFP (PX458) plasmid with the guide sequence 5’ CACCGCGTTTAGAGCAACAGATCG 3’ and the ssODN sequence: 5’ gggttgcatatttatctttaagGTTCTCCGGAAGATCGTGGATCTGAAAGTGACAAGCATTTTAGAAAGg 3’. pSpCas9(BB)-2A-GFP (PX458) was a gift from Feng Zhang (Addgene plasmid # 48138; http://n2t.net/addgene:48138; RRID:Addgene_48138). HeLa cells stably expressing C2-His6 or C2-Flag were generated by lentiviral transduction, using the pCIG3 vector containing the C2-His6 or C2-Flag sequence. pCIG3 (pCMV-IRES-GFP version 3) was a gift from Felicia Goodrum (Addgene plasmid # 78264; http://n2t.net/addgene:78264; RRID:Addgene_78264). The fibroblasts were incubated at 37°C with 5% CO_2_ and 5% O_2_. All other cell lines were incubated at 37°C with 5% CO_2_ and 21% O_2_.

### Antibodies

The following antibodies were used: anti-FLAG M2 (Sigma, cat# F1804-1MG), anti-Tom20 F-10 (Santa Cruz, cat# sc-17764), anti-C2 C-term (Sigma, cat#HPA027407), anti-cytochrome C 6H2.B4 (bd pharmingen, cat# 556432), anti-OPA1 (BD Bioscience, cat# 612606), anti-β-Tubulin (Sigma, cat# T8328), OMA1 (ProteinTech, cat# 17116-1-AP), anti-C2 (ProteinTech, cat# 19424-1-AP), anti-C2 (ProteinTech, cat# 6602-1-Ig), anti-C10 (Sigma, cat# HPA003440), anti-MIC60 (ProteinTech, cat#430909), anti-MIC19 (Sigma, cat# HPA042935), anti-MIC27 AntI-MIC27 (Sigma, cat# HPA000612), anti-GAPDH (ProteinTech, cat# 60004), anti-HSP90 (ProteinTech, cat# 13171-1-AP), anti-MTHFD2 (ProteinTech, cat# 12270-1-AP), and anti-MTHFD1L (ProteinTech, cat# 16113-1-AP).

### Transmission Electron Microscopy

Mouse fibroblasts were fixed with 4% glutaraldehyde (Electron Microscopy Services) in EM buffer (0.1 N sodium cacodylate at pH 7.4 with 2 mM calcium chloride) for 30 minutes at room temperature and then at 4°C for at least 24 hours. Samples were washed with buffer and treated with 1% osmium tetroxide in 0.1 N cacodylate buffer at pH 7.4 for 1 h on ice, washed and *en bloc* stained with 0.25–1% uranyl acetate in 0.1 N acetate buffer at pH 5.0 overnight at 4°C, dehydrated with a series of graded ethanol and finally embedded in epoxy resins. Ultrathin sections (70 nm) were stained with lead citrate and imaged with a JEOL 1200 EXII Transmission Electron Microscope. Images were evaluated and scored by a blinded experimenter. Mitochondrial were scored as abnormal if they displayed cristae that were ring shaped and detached from the boundary membrane, had a total course of the cristae turned more than 90°, and/or there was extreme variability in mitochondrial diameter in which the region with the thinnest caliber was less than half of the region with greater caliber. Cristae were only scored when the outer and boundary inner membrane surrounding the cristae were sharply in view.

### Oxygen Consumption Rate

Seahorse Extracellular Flux Analyzer XF (Aglient) was used to measure the oxygen consumption rates (OCR) of mouse primary fibroblasts. Fibroblast cells were seeded at 20,000 cells/well in XF96 Cell Culture Microplates coated with 1% gelatin and incubated for 24 hrs at 37°C with 5% CO_2_ in DMEM containing glucose. An hour before the assay, the cells were placed in DMEM lacking bicarbonate and culture plates were moved to a 37°C incubator with atmospheric CO2. The assay measures OCR at basal levels, after addition of 1 μm oligomycin, 2 μm FCCP and 0.5 μm of rotenone and antimycin. Data are represented as means ± standard error of at least 6 biological replicates, normalized by protein concentration as determined by the BCA assay.

### Mass Spectrometry

HEK293^WT^ cells were labelled to >95% incorporation of ^12^C_6_,^15^N_2_ L-Lysine and ^13^C_6_,^15^N_4_ L-Arginine (Cambridge Isotope Laboratories) in SILAC DMEM media lacking L-Lysine and L-Arginine (Thermo Fischer). Mitochondria were isolated from both heavy labeled HEK293^WT^ and light labeled HEK293^DKO^ cells. Protein concentration was determined by BCA assay and heavy and light mitochondria were mixed 1:1. The mixed mitochondria were solubilized in 1% digitonin and complexes were separated on a BN-PAGE gels with bovine heart mitochondria used as a molecular weight standard. Lanes were cut into ten gel slices and digested with trypsin. Extracted peptides were desalted and used for LC-MS/MS data acquisition on an Orbitrap Luminos mass spectrometer (Thermo Fisher Scientific) coupled with a 3000 Ultimate high-pressure liquid chromatography instrument (Thermo Fisher Scientific). Peptides were separated on an ES802 column (Thermo Fisher Scientific) with mobile phase B increasing from 2 to 27% over 60 min. The LC-MS/MS data were acquired in data-dependent mode. The resolution of the survey scan (300–1600 m/z) was set at 60k at m/z 400 with a target value of 10 × 106 ions. Collision-induced dissociation was performed on the top ten most abundant precursor ions with an isolation window of 2.0 Da. Database search and H/L ratio calculation were performed using MaxQuant against Sprot Human database (Tyanova et al., 2016). Oxidation (M), Label:13C(6)15N(2) (K), and Label:13C(6)15N(4) (R) were included as variable modifications in the database search. Identified protein groups tagged as reverse, contaminants, or identified only by modified peptide were filtered out using the MaxQuant companion program, Perseus. For the HEK293^WT^ condition intensities were normalized for each protein group across the gel slices by dividing protein intensities in each gel slice by the sum of the intensities from all of the gel slices. Intensities for the HEK293^DKO^ values were calculated by multiplying the relative wildtype value for each protein group in each gel slice by the L/H ratio quantified for each slice. When a ratio was missing the WT value was multiplied by 1.

### Transfection, Immunocytochemistry, and Confocal Microscopy

Cells were plated on 8-well chambered slides (Ibidi) at a seeding density of 25,000 cells/well overnight. Where indicated cells were transfected with plasmids using FugeneHD (Promega). C10 (del ΔH) was produced by cloning a gene block (IDT) comprising a human codon optimized C10 Δ(45 - 69) into a YFP-N1 vector (Clontech) digested with NotI and BamHI to replace the YFP insert, using HiFi NEB-Builder (NEB). Other plasmids were cloned as described previously (Huang et al., 2018). For siRNA knockdown experiments, cells were plated on day 1, transfected with siRNA on day 2, transfected again on day 4 and fixed on day 6. OPA1 was targeted using SMARTpool: On-TARGETplus (Dharmacon, Cat# L-005273-00-0005) and Mic60/IMMT was targeted using siGENOME Human IMMT (10989) siRNA – SMARTpool (Dharmacon, Cat# M-019832-01-0005), using RNAiMAX (Thermo Fischer). Cells were fixed for 10 min with 4% paraformaldehyde (Electron Microscopy Services) in PBS 1X. Cells were then permeabilized with 0.25% Triton X-100, blocked with 1% BSA for at least 30 minutes, and then incubated in primary antibody for 2 hrs to overnight in PBS at 4°C. Cells were washed with PBS three times and then incubated at room temperature for 1 hour with Alexa Fluro goat secondary antibodies (488, 555, 594 or 647) at 1:1000 dilution in 1% BSA. Confocal microscopy was performed on Fluoview3000 (Olympus) or Airy LSM 880 (Zeiss). The antibodies used for immunocytochemistry were anti-FLAG M2 (Sigma cat# F1804-1MG), anti-Tom20 F-10 (Santa Cruz, cat# sc-17764), anti-C2 C-term (Sigma, Cat#HPA027407), anti-HA (3F10) rat monoclonal (Sigma, 11867423001), and anti-cytochrome C 6H2.B4 (BD Pharmingen, cat# 556432).

Fragmentation of the mitochondria in HeLa cells stained with Mitotracker Red (Thermo Fischer) or immunostained for cytochrome *c* or Tom20 was scored to fall in into one of three categories: tubular, short-tubes, and fragmented mitochondrial network. Cells with and without transfection were determined through anti-Flag M2 (Sigma) immunocytochemistry staining. To determine number of cells with greater than 2 foci, HeLa cells stained with Tom20 (Santa Cruz) and C2 (Sigma) were imaged with confocal microscopy. Experimenter counted number of cells in each condition that contained greater than 2 foci visible by wide-field fluorescent microscopy using a 60X objective. To determine C2 foci count and intensity in HeLa cells, Imaris software was used to identify and calculate fluorescent intensity from z-stack confocal images. The “Spots” detection tool in Imaris was used to identify foci from C2 (Sigma) immunostaining. For colocalization of C10 and C2 foci, z-stack confocal images were analyzed in Imaris. The “Spots” tool was used to detect C2 and C10 foci. Then, the total number of C10 foci that colocalized with any C2 focus was calculated. Two foci are considered colocalized if their centers were within 0.01 mm.

### Echocardiography

Mice were lightly anesthetized with isoflurane during exams and placed in the supine position over a heated platform with electrocardiography (ECG)leads and a rectal temperature probe. Heart images were acquired using the Vevo2100 ultrasound system (VisualSonics, Toronto, Canada) with a 30 MHz ultrasound probe (VisualSonics, MS-400 transducer). Measurements were made from standard 2D and M-mode images from the parasternal long axis and mid-papillary short axis views of the LV.

### Histology Analysis

Hearts were harvested from mice anesthetized with isoflurane. Harvested hearts were cut into three segments with a razor blade. The apex and base were snap frozen and a mid-section was fixed in 4% paraformaldehyde (Electron Microscopy Services) for 24 hrs at 4°C, washed with PBS, dehydrated, and embedded in paraffin (through HistoServ), sliced into 5 μm sections and stained with H&E and Masson trichrome (through HistoServ). Sections were imaged with a wide-field microscope (Zeiss).

For immunohistochemistry, deparaffinized cardiac sections were treated in blocking buffer (5% goat serum in 0.3% Triton X-100, 0.02% NaN_2_). The sections were then incubated in primary antibody anti-C10 (1:1000, Sigma-Aldrich, HPA003440) and anti-Tom20 (Santa Cruz sc-17764 1:100 dilution) in PBS overnight at 4°C. After washes in PBS, the sections were incubated with secondary antibodies (Alexa Fluro-labelled goat antibodies) in blocking buffer for 1hr at room temperature. After immunostaining, the tissue was mounted with mounting agent (KPL 71-00-16) and imaged with constant microscopy settings on Fluoview3000 confocal microscope (Olympus) with 40X silicon oil immersion lens. The antibodies used are anti-Tom20 F-10 (Santa Cruz, cat# sc-17764) and anti-C10 (Sigma, cat# HPA003440).

### Immunoblotting

Lysates from HEK293, HeLa, and fibroblast cells were processed as described previously (Huang et al., 2018). Mouse heart and skeletal muscle were lysed in RIPA buffer and a buffer contains 20mM Tris pH 7.8, 137mM NaCl, 2.7mM KCl, 1mM MgCl2, 1% TX-100, 10% glycerol, 1mM EDTA and 1mM DTT respectively with 1% proteinase inhibitor. Lysates were sonicated by a Vibra-Cell Ultrasonic Disruptor for 15s, 4 times at an output level of 20. Protein concentration was determined by the BCA assay. Lysates were separated on SDS-PAGE gels and analyzed by immunoblotting. OMA1 produced S-OPA1 cleavage products were calculated by measuring the maximum intensity of each of the five bands in a linescan of the optical density of the five OPA1 bands on the blot using the software program FIJI. The peaks of c and e bands were summed and divided by sum of the five bands (a - e) to obtain percentage OMA1-generated S-OPA1 of total OPA1.

### RNA Expression Studies

For each for the following groups, RNA was extracted from hearts of four mice: WT younger, WT older, C10^S59L/+^ younger, C10^S59L/+^ older, and C2/C10 DKO mice. RNA expression was measured using the Clariom_S_Mouse microarray (Affymetrix). Data was analyzed using the Transcriptome Analysis Console (TAC) Software (version 4.0) on the default settings (Affymetrix). Specifically, data was summarized using the Gene Level – SST-RMA model. Differentially expressed genes (DEGs) were required to have gene-level fold change < −2 or > 2 and gene level P-Value < 0.05 measured using the ebayes Anova method. Additionally, expression levels of a pre-specified list of 24 individual genes consisting of C2, myc, and 22 genes previously identified as mt-ISR related DEGs in an independent C10^S59L/+^ mouse line were extracted for each sample and analyzed nominally using one-way ANOVA with Sidak’s multiple comparison test applied for each gene.

### Statistical Analysis

For all statistical analyses with 2 samples, Student’s *t*-tests (2-tailed) were performed in Excel (Microsoft) or Prism (Graphpad). For analyses comparing more than 2 samples one-way ANOVA with Sidak’s multiple comparison test was performed in Prism (Graphpad).

## Acknowledgements

We thank Maric Dragan PhD and the NINDS Flow Cytometry Core Facility for technical assistance with FACS experiments. We thank Virginia Crocker and the NINDS EM Facility for technical assistance with transmission electron microscopy. We thank Carolyn Smith PhD and the NINDS Light Microscopy Facility for technical assistance with confocal microscopy. We thank Abdel Elkahloun PhD and the NHGRI/DIR Microarray Core for technical assistance with RNA expression studies. We thank Yan Li at the NINDS Protein/peptide Sequencing Facility for the SILAC-based proteomics analysis. We thank Richard Youle PhD for critical reading of the manuscript and insightful comments. This work was supported by the Intramural Research Program of the NINDS, National Institutes of Health.

## Conflict of Interest Statement

None declared.

## Supplemental Methods

### Antibodies

The following antibodies were used: MTHFD2 (Proteintech, cat# 122701-1-AP), MTHFD2 (Abcam, # ab1151447), Mic25 (Proteintech, cat# 20639-1-AP, Drp1 (BD Biosciences, cat# 611113), MFN2 (kind gift from Richard Youle [NINDS, Bethesda]), PGAM5 (Santa Cruz, cat# SC-515880, YME1L1 (Proteintech, cat# 11510-1-AP, SLP2 (Proteintech, cat# 10348-1-AP, PARL (Proteintech, cat# 26679-1-AP).

### Measurement of membrane potential in primary fibroblasts by TMRM

Membrane potential of primary fibroblasts was measured using TMRM using an automated high content imaging system as described previously (Ashley et al., 2005). For live cell staining, 500 μM PicoGreen and 500 μM TMRM stocks were diluted in cell culture media 1:333 (3µl/ml) and 1:2000 respectively. 100µl of staining solution was added to each well of a 96-well plate. Fibroblasts were incubated in the staining solution for 30 mins at 37°C in a humidified incubator with 5% CO_2_. The staining solution was removed and replaced with 150µl pre-warmed reduced serum Opti-MEM media for imaging. Image stacks were analysed by an in-house written protocol using the IN Cell Developer software (GE Healthcare) (Diot et al., 2015). A series of target sets were created by the segmentation of cellular components. These target sets were then linked together in order to create a set of cells with all the labelled organelles in which various measurements were carried out on a cell-by-cell bases. The cell body, the nucleus and mtDNA were segmented in the green channel (PicoGreen). Mitochondria were segmented in the red channel (TMRM).

## Supplemental Figure Legends

**Supplemental Figure 1.**
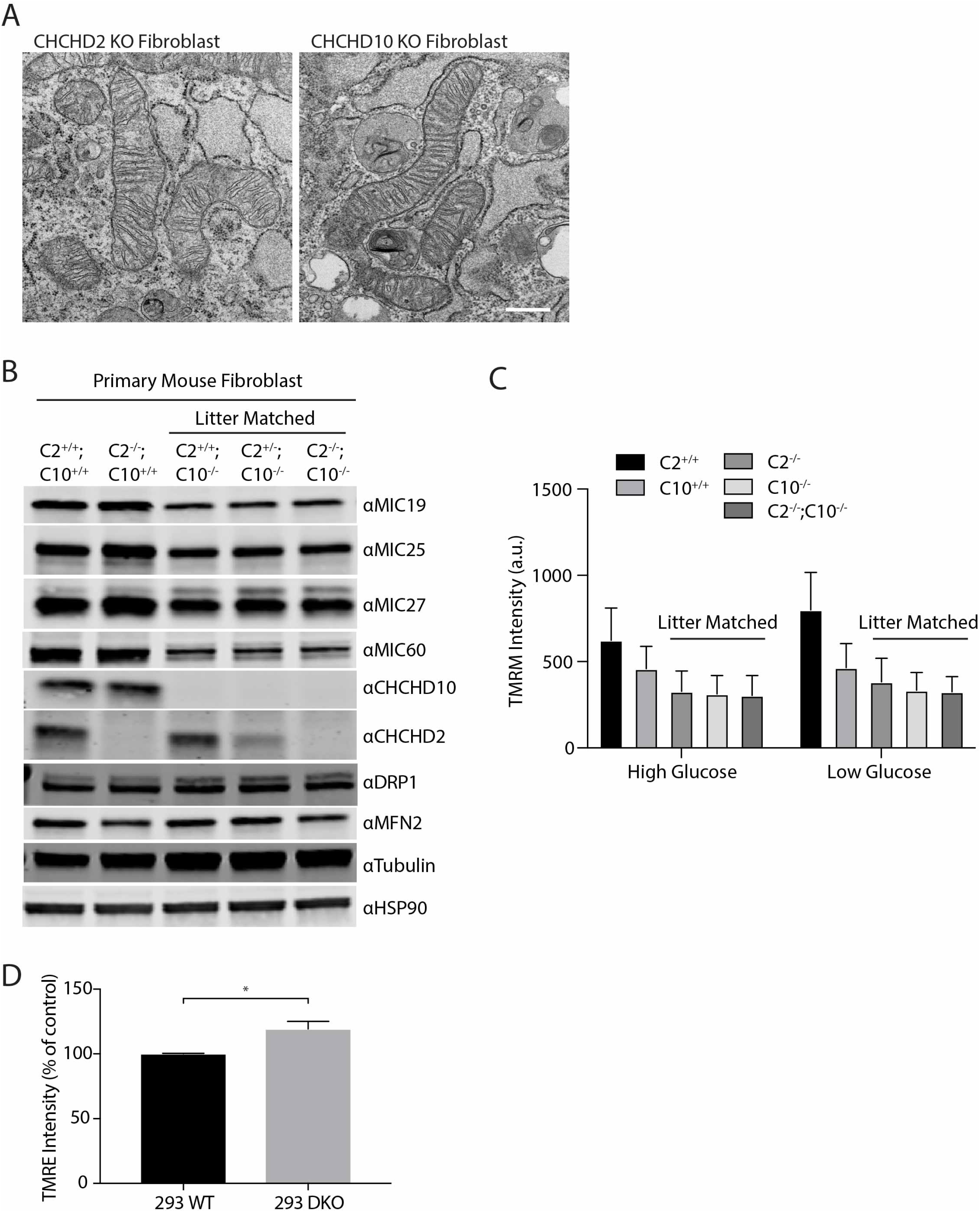
MICOS complex subunits and membrane potential and normal PGAM5 processing in C2/C10 DKO cells. (A) Representative TEM images of C2 KO and C10 KO primary mouse fibroblasts. (B) Immunoblot of MICOS subunits in primary fibroblasts. (C) TMRM intensity of primary fibroblasts measured by automated high content microscopy. (D) Relative TMRE fluorescence in HEK293 WT and DKO cells measured by flow cytometry. N = 3. “*” indicates p-value < 0.05. Lines from litter matched C2^+/+^C10^-/-^, C2^+/-^C10^-/-^, and C2^-/-^C10^-/-^ mice are indicated.

**Supplemental Figure 2.**
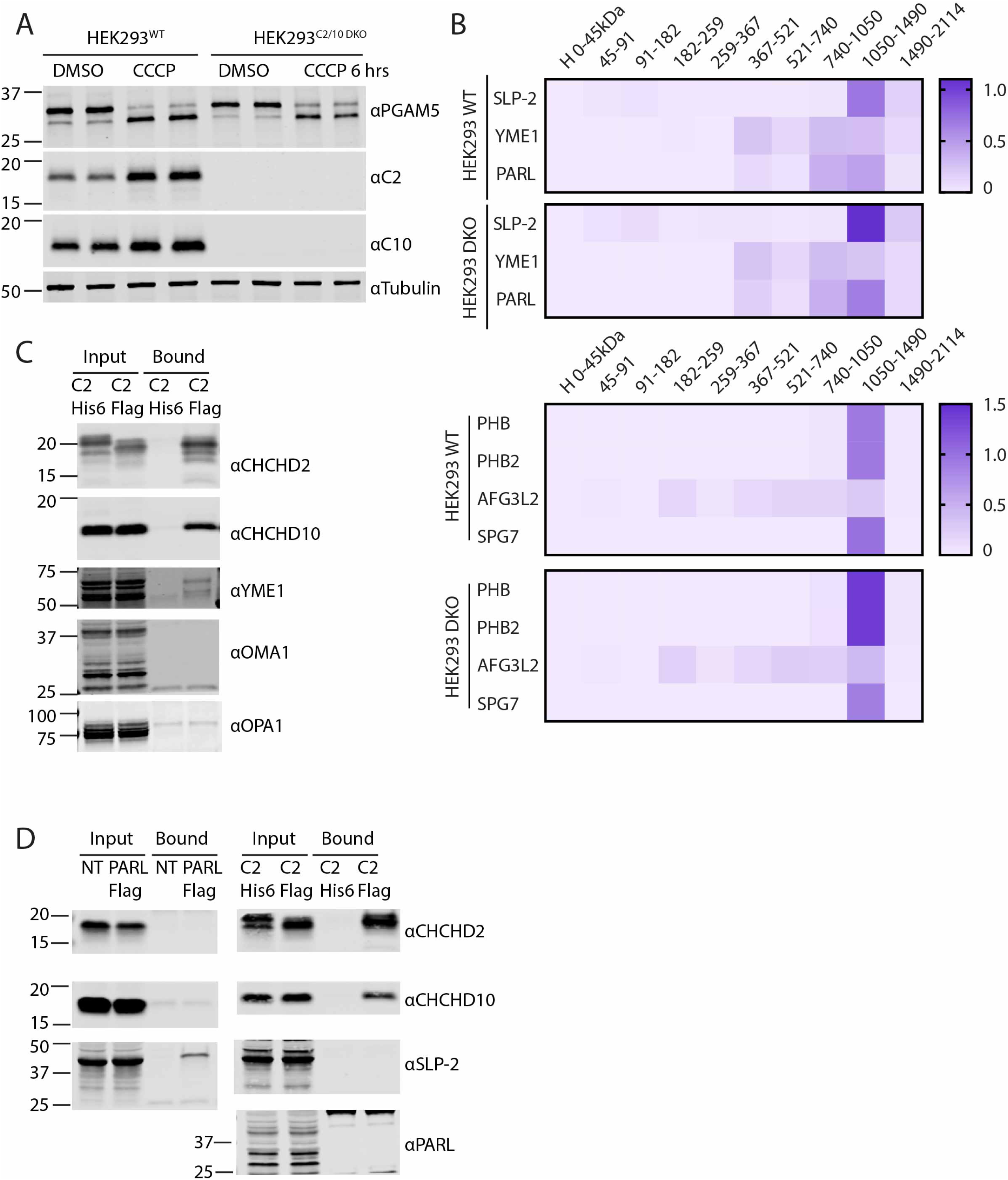
Interactions between C2/C10 and OPA1, OMA1, and SPY complexes. (A) Immunoblot of PGAM5 processing in HEK293 WT and C2/C10 DKO cells. (B) Heat map depicts relative abundance of SPY and PHB complex subunits in SILAC labelled WT and DKO HEK293 mitochondria, solubilized in 1% digitonin and separated by BN-PAGE prior to detection by quantitative mass spectrometry. Columns in the heat map represent gel slices at the indicated position. Rows indicate proteins detected. (C) Immunoblot of unbound and bound fractions following immunoprecipitation with anti-Flag antibodies from HeLa cells stably expressing C2-His6 (as a control) or C2-Flag. (D, left) Immunoblot of unbound and bound fractions following immunoprecipitation with anti-Flag antibodies from HeLa^PARL KO^ cells transiently expressing PARL-Flag or left untransfected. (D, right) Immunoblot of unbound and bound fractions following immunoprecipitation with anti-Flag antibodies from HeLa cells stably expressing C2-His6 (as a control) or C2-Flag.

**Supplemental Figure 3.**
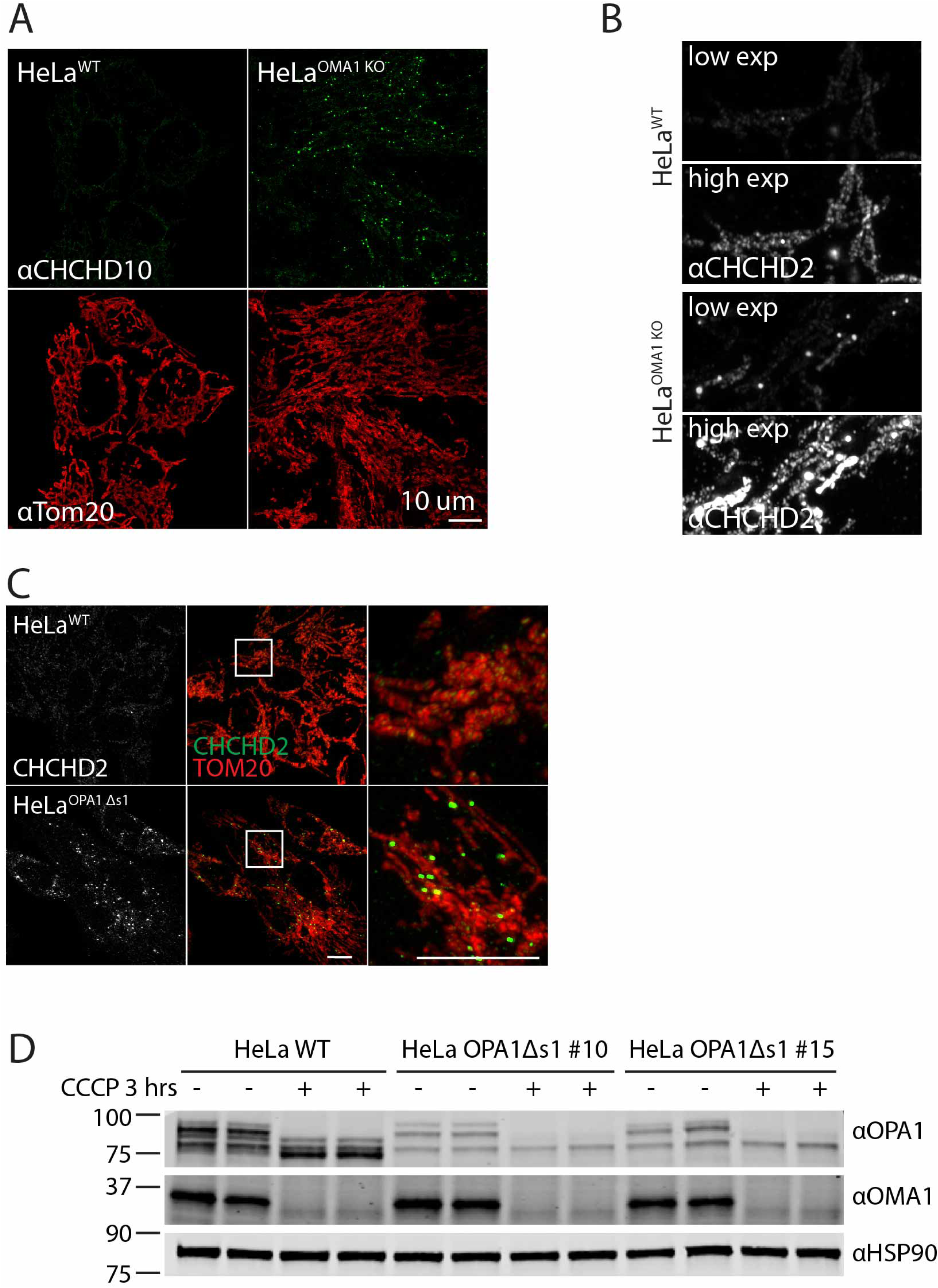
C2/C10 foci form due to altered OPA1 processing. (A) Airyscan confocal image of WT or OMA1 KO HeLa cells immunostained for C10 or Tom20. Scale bar = 10 μm. (B) Airyscan confocal images of WT and OMA1 KO HeLa cells immunostained for CHCHD2 acquired at settings optimized for intense foci in OMA1 KO HeLa cells (low exposure) and WT HeLa cells (high exposure). Scale bar = 5 μm. (C) Representative Airyscan confocal images of WT or OPA1Δs1 HeLa cells immunostained for CHCHD2 (green) and Tom20 (red). (D) Immunoblot of HeLa^WT^ and two HeLa^OPA1Δs1^ clones (#10 and #15) treated with DMSO or 10 μM CCCP for 3 hrs. HSP90 served as a loading control.

**Supplemental Figure 4.**
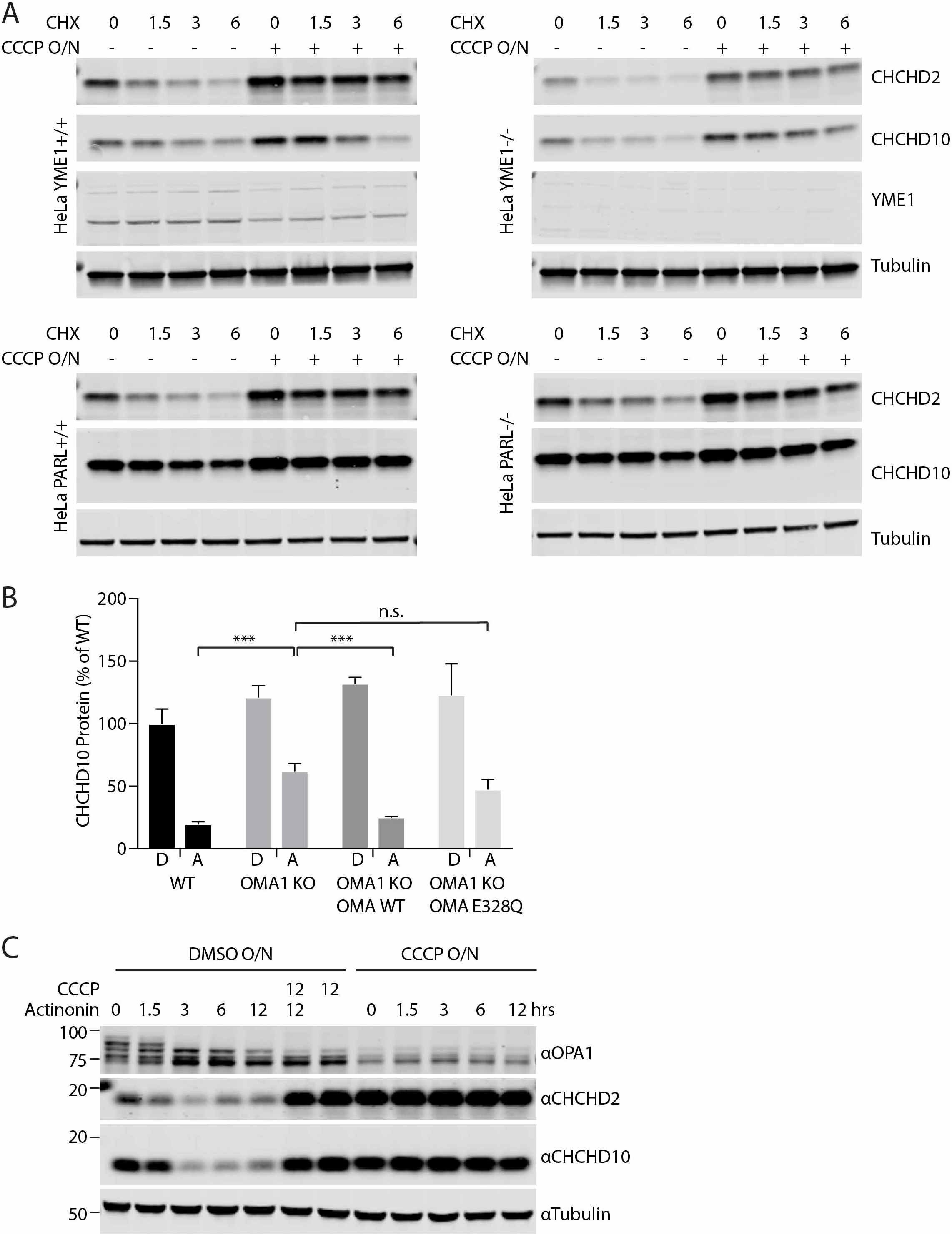
C2/C10 degraded following OMA1 activation by actinonin and basal degradation of C2/C10 is independent of YME1 and PARL. (A) Immunoblot of YME1 KO, PARL KO, and WT HeLa cells treated with DMSO or 10 μM CCCP overnight (O/N) and then treated with 100 μM cycloheximide (CHX) for the indicated number of hours. GAPDH or Tubulin served as controls where indicated. (B) Quantification of the C10 levels from (Figure 5H). N = 3 biological replicates per sample. (C) Immunoblot of WT HeLa cells treated with DMSO or 10 μM CCCP overnight (O/N), followed by the addition of actinonin, CCCP, or 125 μM actininon + 10 μM CCCP for the indicated number of hours. Tubulin served as a loading control.

**Supplemental Figure 5.**
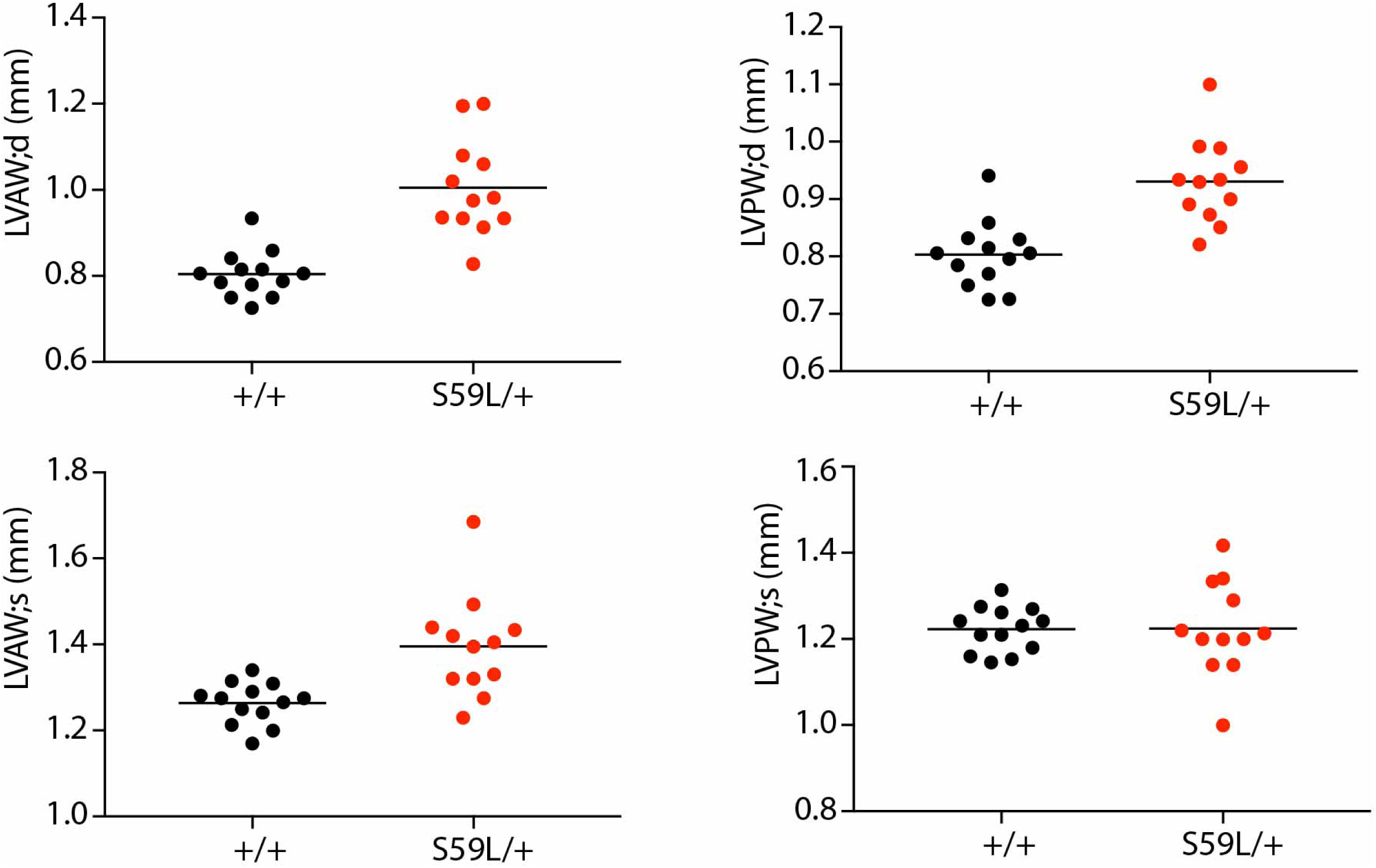
Cardiomyopathy of C10^S59L/+^ mice measured by echocardiography. Echocardiographic measurements of anterior and posterior left ventricular wall widths in diastole (LVWA;d and LVWA;d) and systole (LVWA;s and LVWA;s).

**Supplemental Figure 6.**
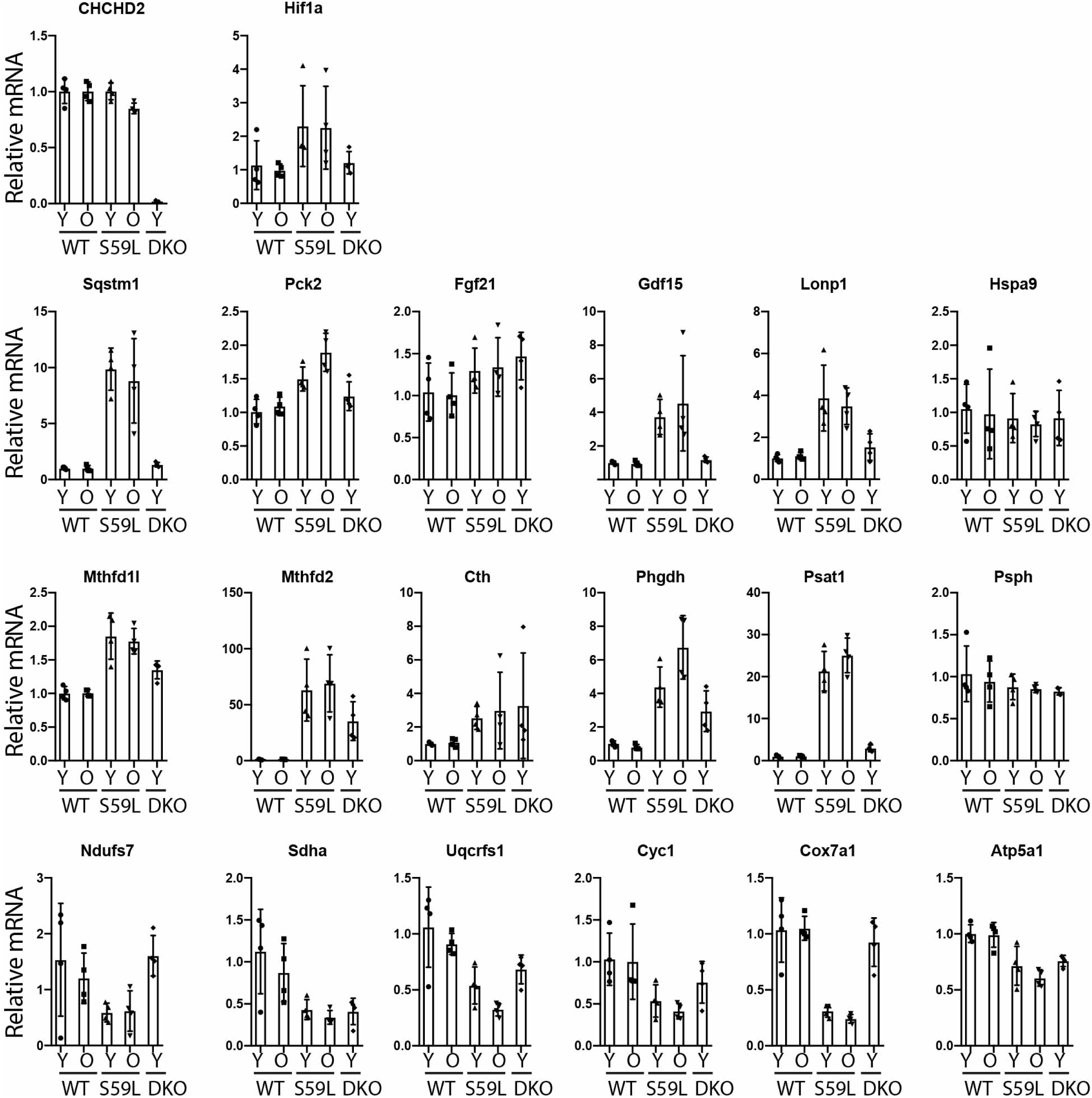
Expression of pre-specified genes associated with mitochondrial integrated stress response and previously reported to be differentially regulated in C10^S59L/+^ mouse hearts. Relative transcript levels pre-specified genes associated with mitochondrial integrated stress response in heart extracts of younger WT, C10^S59L/+^ KI, and C2/C10 DKO mice (9 - 13 weeks), and older WT and C10^S59L/+^ KI (22 - 37 weeks) mice measured by microarray and normalized to average of younger WT mice.

**Supplemental Figure 7.**
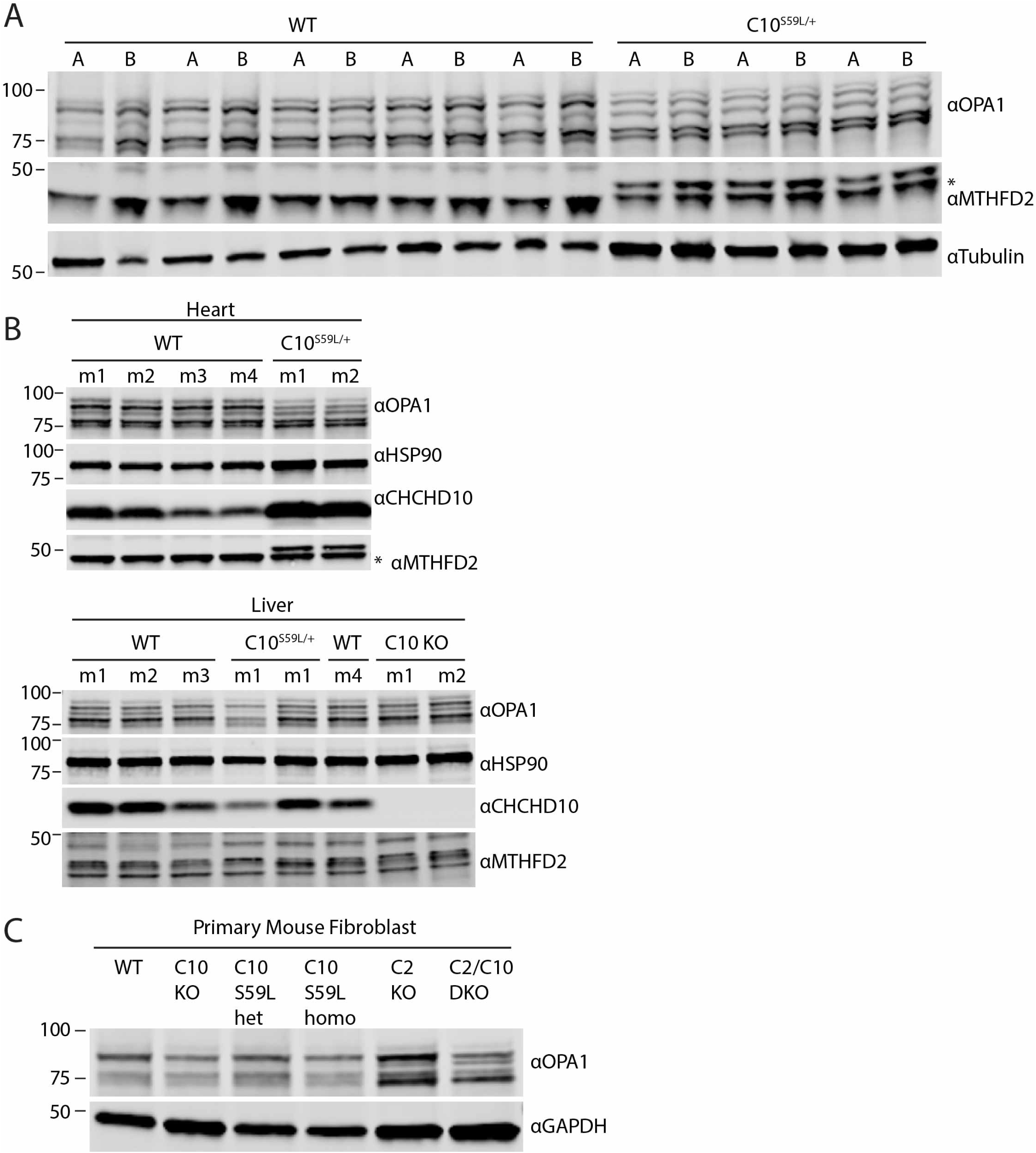
OPA1 processing in tissues from C10^S59L/+^ knock-in mice. (A) Immunoblot of lysates from apex (A) or base (B) of hearts of wildtype or C10^S59L/+^ knock-in mice. Tubulin served as a loading control. (B) Immunoblot of lysates from the heart, skeletal muscle, or liver of wildtype or C10^S59L/+^ knock-in mice. HSP90 served as a loading control. Lysates from individual mice in each condition are indicated (m1 – m4). (C) Immunoblot of primary fibroblasts lysates from WT, C10 KO (C10 KO), C10^S59L/+^, C10^S59L/S59L^, C2 KO (C2 KO), and C2/C10 DKO (C2/C10 DKO) mice. GAPDH served as a loading control. * indicates non-specific band.

**Supplemental Figure 8.**
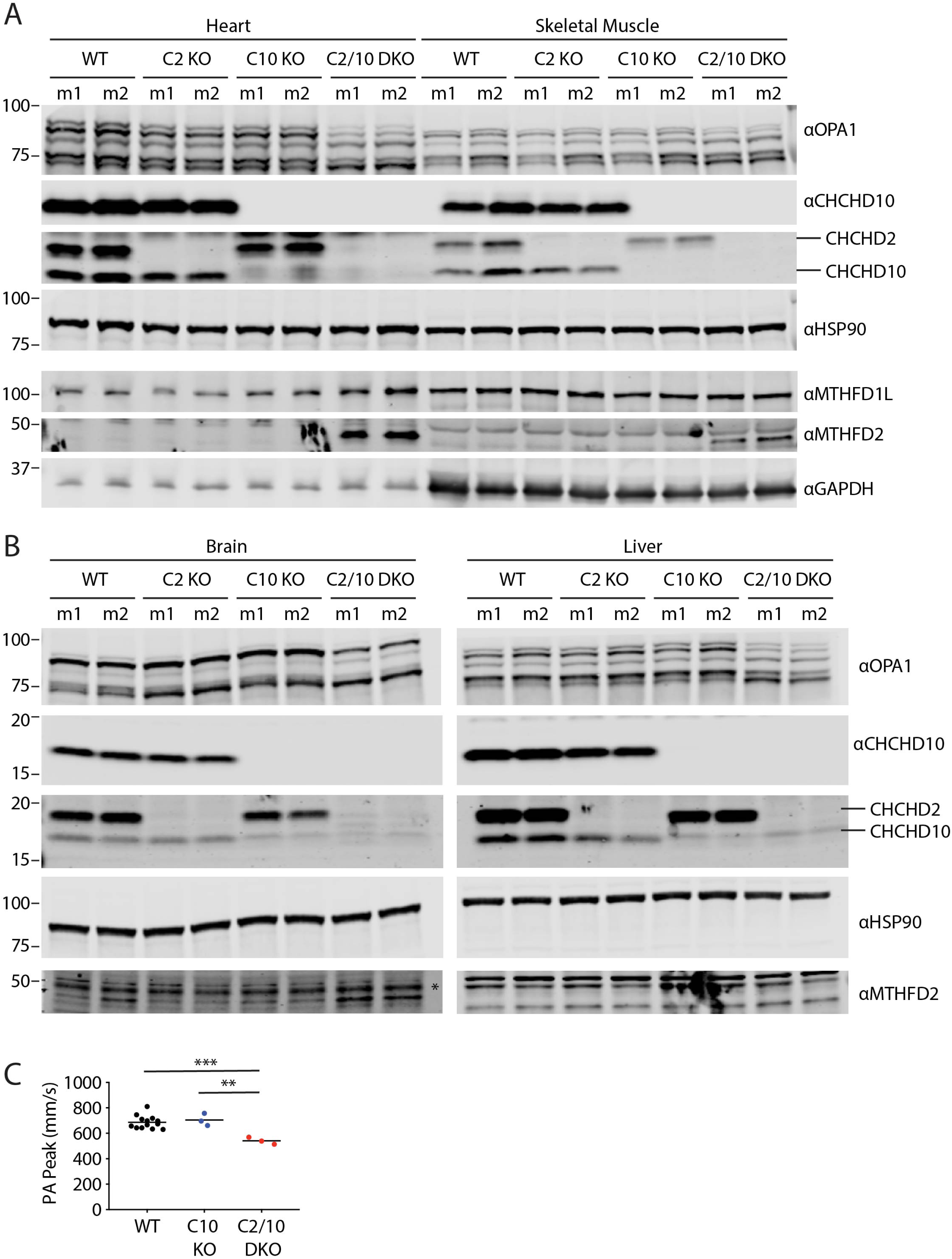
OPA1 processing in tissues from C2/C10 DKO mice. (A and B) Immunoblot of lysates from the heart and skeletal muscle (A) and brain and liver (B) of wildtype or C2/C10 single or double KO mice. HSP90 served as a loading control. Lysates from individual mice in each condition are indicated (m1 – m2). (C) Graph depicts peak pulmonary artery (PA) velocities from WT, C10 KO, and C2/C10 DKO mice measured by doppler echocardiography.

Table S1. LC-MS BN-PAGE complexomic profiling.

Table S2. Mutant C10 and C2/C10 DKO echocardiography.

Table S3. Mutant C10 and C2/C10 RNA expression studies.

